# Exploring the therapeutic potential of a novel series of imidazothiadiazoles targeting focal adhesion kinase (FAK) for pancreatic cancer treatment: Synthesis, mechanistic Insights and promising antitumor and safety profile

**DOI:** 10.1101/2024.05.27.596063

**Authors:** Daniela Carbone, Camilla Pecoraro, Fabio Scianò, Francesca Terrana, Geng Xu, Cecilia Bergonzini, Margot S F Roeten, Stella Cascioferro, Girolamo Cirrincione, Patrizia Diana, Elisa Giovannetti, Barbara Parrino

## Abstract

Focal Adhesion Kinase (FAK) is a non-receptor protein tyrosine kinase that plays a crucial role in various oncogenic processes related to cell adhesion, migration, proliferation, and survival. The strategic targeting of FAK represents a burgeoning approach to address resistant tumors, such as pancreatic ductal adenocarcinoma (PDAC).

Herein, we report a new series of twenty imidazo[2,1-*b*][1,3,4]thiadiazole derivatives assayed for their antiproliferative activity against the National Cancer Institute (NCI-60) panel and a wide panel of PDAC models. Lead compound **10l** exhibited effective antiproliferative activity against immortalized (SUIT-2, CAPAN-1, PANC-1, PATU-T, BxPC-3), primary (PDAC-3) and gemcitabine-resistant clone (PANC-1-GR) PDAC cells, eliciting IC_50_ values in the low micromolar range (1.04-3.44 µM), associated with a significant reduction in cell-migration and spheroid shrinkage *in vitro*. High-throughput kinase arrays revealed a significant inhibition of the FAK signalling network, associated to induction of cell cycle arrest in G2/M phase, suppression of tumor cell invasion and apoptosis induction. The low selectivity index/toxicity prompted studies using PDAC mouse xenografts, demonstrating significant inhibition of tumor growth and safety. In conclusion, compound **10l** displayed antitumor activity and safety in both *in vitro* and *in vivo* models, emerging as a highly promising lead for the development of FAK inhibitors in PDAC.

## INTRODUCTION

Despite advancements in anticancer research, the persistently high global cancer mortality rate, currently at 10 million deaths per year, is expected to increase to 12 million by 2030 due to a projected rise in life expectancy [1]. Pancreatic ductal adenocarcinoma (PDAC), comprising more than 90% of pancreatic cancer cases, stands out as one of the deadliest tumors, with a 5-year overall survival rate of merely 13% [2]. It ranks as the seventh leading cause of cancer-related deaths globally, [3] but it is predicted to become the second cause of cancer-related deaths by 2030 due to rising incidence (1.1% annual growth) and to the very limited therapeutic options [4].

Currently, surgical resection remains the sole potential curative option for PDAC; however, this malignancy presents a considerable diagnostic challenge, often being diagnosed at advanced stages with local invasion or distant metastasis [5]. This late presentation is attributed to various factors, such as the non-specific symptoms associated with the disease and the proximity of vital blood vessels, which are susceptible to tumor invasion. Consequently, the vast majority of pancreatic tumors (around 80-85%) are deemed unresectable upon diagnosis [6]. The first-line treatment for locally-advanced or metastatic PDAC patients consists of cytotoxic chemotherapy, primarily FOLFIRINOX and gemcitabine-based regimens [7–9]. Indeed promising immunotherapeutic approaches that have shown success in various solid tumors, such as melanoma or renal cell carcinoma, are not applicable in the context of PDAC due to its nonimmunogenic, immune-suppressive, and therapy-resistant microenvironment [10]. Given this context, there is an urgent need for novel therapeutic approaches to effectively combat this malignancy [11].

Recently, we reported the synthesis and the remarkable antiproliferative activity against the PDAC cell lines SUIT-2, CAPAN-1 and PANC-1, of a novel series of imidazo[2,1-*b*][1,3,4]thiadiazole compounds of type **1** (Figure 1) [12].

**Figure 1.**
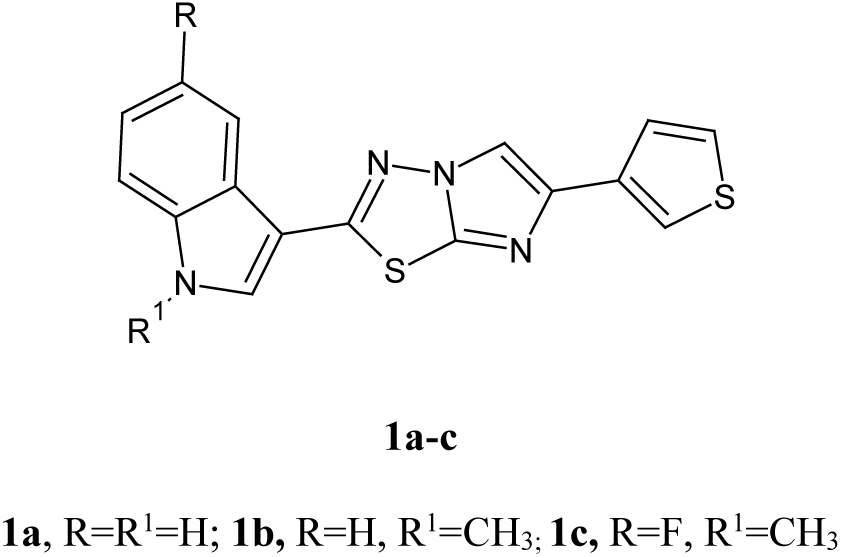
Chemical structure of imidazo[2,1-*b*][1,3,4]thiadiazoles **1a-c**.

In particular, derivatives **1a-c**, bearing the imidazothiadiazole central core substituted in position 2 with an indole moiety and in position 6 with a thiophene ring, elicited GI_50_ values ranging from sub-micromolar to micromolar levels (0.85-2.37 µM). A marked anti-migratory activity of compounds **1** was observed in several PDAC preclinical models and mechanistic studies, employing both kinase arrays and a specific ELISA assay, revealed that these compounds exert their action by inhibiting the phosphorylation of PTK2/FAK (protein tyrosine kinase 2/Focal adhesion kinase) [13].

By binding to the death domain of receptor-interacting protein (RIP), FAK plays a key role in regulating adhesion and cell migration dynamics, as well as promoting cell cycle progression [14–16]. Furthermore, FAK aids in cell survival under adverse growth conditions by enhancing the proteasomal degradation of the tumor suppressor p53 [17,18].

Prior research has demonstrated the anticancer properties of 1,3,4-thiadiazole derivatives containing the 1,4-benzodioxane scaffold, with their primary efficacy attributed to the inhibition of FAK [19]. Notably, the presence of the 1,4-benzodioxine moiety, widely used for its versatility when designing molecules with various biological activities, was identified as crucial for FAK inhibition in these compounds [20–22]. Analogously, the furan ring was identified as promising scaffold for the development of novel potent dual inhibitors targeting both EGFR and FAK [23,24].

Building upon the encouraging results obtained for the compounds of type **1** together and on the promising anti-FAK properties reported for compounds bearing the 1,4-benzodioxine and the furan ring, as well as leveraging our knowledge in nitrogen heterocyclic systems with anticancer properties [25–30], we synthesized a new series of imidazothiadiazole compounds. In particular, the thiophene ring in position 6 was replaced with either a benzodioxane or a furan moiety, in order to obtain potent anticancer agents with an increased capacity to inhibit FAK.

### Chemistry

Twenty new imidazothiadiazole derivatives **10a-t** were efficiently synthesized following the synthetic route described in Scheme 1. In particular, the carbonitriles **3b-e,** not commercially available, were prepared from appropriate 1H-indoles **2** with chlorosulfonyl isocyanate (CSI) in anhydrous acetonitrile under stirring at 0 °C as previously described [31].

**Scheme 1.**
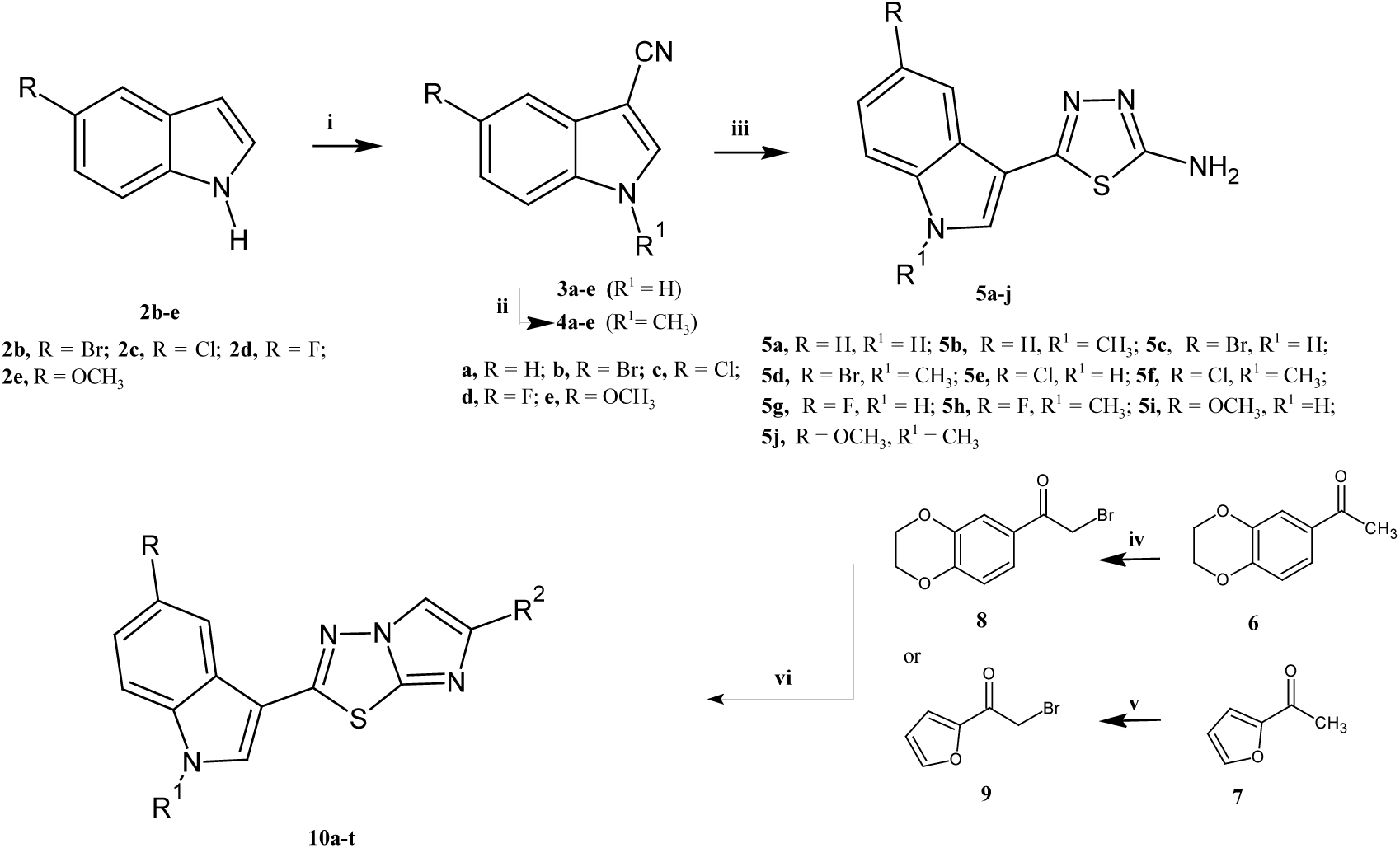
Synthesis of imidazo [2,1-*b*][1,3,4]thiadiazole derivatives **10a-t**. Reagents and conditions: i) CH_3_CN, CSI, 0 °C, 2 h, then DMF, 0 °C, 1.5 h (98-100%); ii) DMF, (CH_3_O)_2_CO, K_2_CO_3_, 130 °C, 3.5 h (98-100%); iii) trifluoroacetic acid, thiosemicarbazide, 60 °C, 3.5 h (98-100%); iv) pyridinium bromide-perbromide in anhydrous MeOH, anhydrous DCM, r.t., 24 h (70%); v) Br_2_, ethyl ether, 24 h, (75%); vi) anhydrous ethanol, reflux, 24 h (47-83%).

The methylation of derivatives **3,** performed with dimethyl carbonate in anhydrous DMF under reflux at 130 °C, gave the corresponding 1-methyl-1H-indole-3-carbonitriles **4**. The thiadiazole intermediates **5a-j** were obtained in excellent yields by the reaction of derivatives **3a-e** or **4a-e** with thiosemicarbazide in trifluoroacetic acid (TFA) at 60 °C for 3 hours. Ultimately, by refluxing in anhydrous ethanol the thiadiazolamines **5** with the suitable α-bromoacetyl derivatives **8** or **9**, the desired compounds **10a-t** were obtained in good yields, ranging from 47% to 78% (Table 1).

**Table 1.** New imidazothiadiazole compounds 10a-t.

| Cpd | R | R<sup>1</sup> | R<sup>2</sup> | Cpd | R | R<sup>1</sup> | R<sup>2</sup> |
| --- | --- | --- | --- | --- | --- | --- | --- |
| **10a** | H | H | 2,3-dihydro-benzo[1,4]dioxin-6-yl | **10k** | H | H | furan-2-yl |
| **10b** | H | CH<sub>3</sub> | 2,3-dihydro-benzo[1,4]dioxin-6-yl | **10l** | H | CH<sub>3</sub> | furan-2-yl |
| **10c** | Br | H | 2,3-dihydro-benzo[1,4]dioxin-6-yl | **10m** | Br | H | furan-2-yl |
| **10d** | Br | CH<sub>3</sub> | 2,3-dihydro-benzo[1,4]dioxin-6-yl | **10n** | Br | CH<sub>3</sub> | furan-2-yl |
| **10e** | Cl | H | 2,3-dihydro-benzo[1,4]dioxin-6-yl | **10o** | Cl | H | furan-2-yl |
| **10f** | Cl | CH<sub>3</sub> | 2,3-dihydro-benzo[1,4]dioxin-6-yl | **10p** | Cl | CH<sub>3</sub> | furan-2-yl |
| **10g** | F | H | 2,3-dihydro-benzo[1,4]dioxin-6-yl | **10q** | F | H | furan-2-yl |
| **10h** | F | CH<sub>3</sub> | 2,3-dihydro-benzo[1,4]dioxin-6-yl | **10r** | F | CH<sub>3</sub> | furan-2-yl |
| **10i** | OCH<sub>3</sub> | H | 2,3-dihydro-benzo[1,4]dioxin-6-yl | **10s** | OCH<sub>3</sub> | H | furan-2-yl |
| **10j** | OCH<sub>3</sub> | CH<sub>3</sub> | 2,3-dihydro-benzo[1,4]dioxin-6-yl | **10t** | OCH<sub>3</sub> | CH<sub>3</sub> | furan-2-yl |

The two α-bromoacetyl derivatives **8** or **9,** used in the final reaction, were obtained in good yields following two different procedures. In particular, the 2-bromo-1-(2,3-dihydro-benzo[1,4]dioxin-6-yl)-ethanone **8** was obtained in good yield by treating a solution of 3′,4′-(ethylenedioxy)-acetophenone **6** with pyridinium bromide-perbromide in methanol, whereas, the 2-bromo-1-furan-2-yl-ethanone **9** was prepared through the bromination of 2-acetylfuran **7** with bromine in diethyl ether.

## Material and methods

### Chemistry

The substances necessary to carry out the reported syntheses, both reagents and solvents, have been acquired from Merck, VWR International, Alfa Aesar and Acros Organics. Dichloromethane was anhydrified by using calcium hydride and stored over 4 Å molecular sieves. For the analytical thin-layer chromatography (TLC), silica gel 60 F254 plates (0.25 mm thickness) were used and the developed plates were observed under ultraviolet (UV) light. A Buchi-Tottoly capillary instrument was employed for the melting points evaluation. For recording IR spectra in bromoform, we used a Shimadzu FT / IR 8400S spectrophotometer and reported the peaks in wavenumber (cm^−1^). ^1^H and ^13^C NMR spectra were acquired at 200 and 50 MHz, respectively, through a Bruker Avance II series 200 MHz spectrometer. The purification of the compounds was carried out by using chromatography column with MERK silica gel 230-400 mesh ASTM or with Buchi Sepacore chromatography module (prepacked cartridge reference).

### Synthesis of 1H-indole-3-carbonitriles (3b-e)

Chlorosulfonyl isocyanate (CSI) (0.44 mL, 5.10 mmol) was added dropwise to a solution of the appropriate indole **2** (5.10 mmol) in anhydrous acetonitrile (4.5 mL). After two hours under stirring at 0°, anhydrous dimethylformamide (DMF) (2.8 mL, 36.39 mmol) was slowly added and the reaction mixture was kept at 0 °C for another 1.5 h. The solution was then poured into crushed ice and the solid product obtained was filtered and dried (yields 98-100%). Analytical and spectroscopic data for compounds **3b-e** are in agreement with those previously reported [12,31].

### Synthesis of 1-methylindole-3-carbonitriles (4a-e)

A mixture of the proper 3-cyanoindole **3** (7.03 mmol), anhydrous DMF (10 mL), 3.61 mmol of K_2_CO_3_ and dimethyl carbonate (1.8 mL, 21.4 mmol) was heated under stirring at 130 °C for 3.5 h. After this time, the mixture was cooled at 0-5 °C, and water and ice (25 mL) was slowly added. The extraction of the resulting suspension with diethyl ether (3 × 10 mL), followed by washing the organic phase with brine and drying over Na_2_SO_4,_ allowed to obtain the 3-cyano-1-methylindoles **4** in excellent yields (98-100%). Analytical and spectroscopic data are in accordance to those reported in literature [31].

### Synthesis of 5-(1H-indol-3-yl)-1,3,4-thiadiazol-2-amines (5a-j)

The suitable indole-3-carbonitrile **3a-e,** or the methyl analogue **4a-e** (5 mmol), was treated with thiosemicarbazide (5 mmol) in trifluoroacetic acid (5 mL) under stirring at 60 °C for 3.5 h. After cooling, the mixture was poured into ice and treated with NaHCO_3_ saturated solution up to pH 7. The solid was filtered off and washed with water, cyclohexane and diethyl ether to obtain the desired 5-(1*H*-indol-3-yl)-1,3,4-thiadiazol-2-amines **5a-j** in excellent yields.

Analytical and spectroscopic data for the derivatives **5a-j** are in accordance to those reported in literature [12].

### Symthesis of 2-bromo-1-(2,3-dihydro-1,4-benzodioxin-6-yl)ethanone 8

To a solution of 1-(2,3-dihydro-benzo[1,4]dioxin-6-yl)-ethanone **6** (1.68 mmol) in 3 mL of anhydrous DCM, pyridinium tribromide (3.53 mmol) solubilized in anhydrous methanol (2.8 mL) were slowly added at room temperature and the reaction mixture was stirred for 24 h. The resulting solid precipitate was filtered off and washed with cold ethanol to obtain the pure compound **8** in good yields (70%).

Analytical and spectroscopic data for the derivative **8** are in accordance to those reported in literature [32].

### Symthesis of 2-bromo-1-(furan-3-yl)ethanone 9

To a solution of 2-acetylfuran **7** (20 mmol) in 30 mL of anhydrous diethyl ether, bromine (20 mmol) was added dropwise under stirring at the temperature of 0-5°C (ice bath). The reaction mixture was stirred at room temperature for 24h. After this time, 20 mL of water were added and the obtained solution was extracted with diethyl ether (3 × 10 mL), the combine organic layers were washed with a saturated aqueous NaHCO_3_ and a solution of Na_2_S_2_O_3,_ dried over anhydrous Na_2_SO_4._ The solvent was evaporated under reduced pressure to obtain the crude product **9,** which was purified by column chromatography using DCM/ethyl acetate 9:1 as eluent (75%). Analytical and spectroscopic data for the derivative **9** are in accordance to those reported in literature [33].

### General procedure for the synthesis of 3-[6-(2,3-dihydro-benzo[1,4]dioxin-6-yl)-imidazo[2,1-b][1,3,4]thiadiazol-2-yl]-1H-indole derivatives 10a-j and 3-(6-furan-2-yl-[2,1-b][1,3,4]thiadiazol-2-yl)-1H-indole derivatives 10k-t

5-(1H-Indol-3-yl)-1,3,4-thiadiazol-2-amine **5a-f** (0.92 mmol) and the suitable α-bromoacetyl derivative **3** or **4** (0.92 mmol) were refluxed in 40 mL of anhydrous ethanol for 24 h. For the synthesis of derivatives **10k-t**, 0.13 mL of triethylamine were added to the mixture. After cooling at room temperature the solid was filtered off and washed with cold ethanol, obtaining the imidazothiadiazole **10** as pure compound, except for **10c,f,m-r** which were purified by silica gel column chromatography eluting by dichloromethane - ethyl acetate, 9:1.

### 3-[6-(2,3-Dihydro-benzo[1,4]dioxin-6-yl)-imidazo[2,1-b][1,3,4]thiadiazol-2-yl]-1H-indole (10a)

Yield: 65 %; yellow solid; m.p.: 230 °C; IR (cm^−1^): 3020 (NH); ^1^H NMR (200 MHz, DMSO-*d_6_*) δ: 4.28 (4H, s, CH_2_), 6.94 (1H, d, *J* = 8.2 Hz, Ar), 7.27-7.56 (5H, m, Ar), 8.15 (1H, d, *J* = 5.4 Hz, Ar), 8.40 (1H, s, Ar), 8.71 (1H, s, Ar), 12.21 (1H, s, NH).^13^C NMR (50MHz, DMSO-*d_6_*)δ: 64.6 (t), 64.7 (t), 106.6 (s), 106.9 (s), 113.1 (d), 113.8 (d), 115.1 (d), 117.9 (d), 118.3 (d), 119.7 (d), 120.8 (d), 122.1 (d), 123.8 (d), 124.2 (s), 126.1 (s), 126.7 (s), 137.2 (s), 142.9 (s), 143.7 (s), 144.1 (s). Anal. calculated for C_20_H_14_N_4_O_2_S (MW 374.42): C, 64.16; H, 3.77; N, 14.96%. Found: C, 64.32; H, 3.81; N, 15.01%.

### 3-[6-(2,3-Dihydro-benzo[1,4]dioxin-6-yl)-imidazo[2,1-b][1,3,4]thiadiazol-2-yl]-1-methyl-1H-indole (10b)

Yield: 67 %; light brown solid; m.p.: 230 °C; ^1^H NMR (200 MHz, DMSO-*d_6_*) δ: 3.90 (3H, s, CH_3_), 4.29 (4H, s, CH_2_), 6.94 (1H, d, *J* = 8.3 Hz, Ar), 7.31-7.38 (4H, m, Ar), 7.62 (1H, d, *J* = 7.4 Hz, Ar), 8.14 (1H, d, *J* = 6.6, Ar), 8.40 (1H, s, Ar), 8.69 (1H, s, Ar). ^13^C NMR (50 MHz, DMSO-*d_6_*) δ: 33.6 (q), 64.4 (t), 65.1 (t), 105.6 (d), 105.9 (s), 106.7 (s), 110.2 (d), 111.5 (d), 112.1 (d), 121.0 (d), 122.3 (d), 123.7 (d), 124.5 (s), 126.2 (s), 127.9 (s), 133.5 (d), 137.6 (s), 137.7 (d), 142.4 (s), 149.7 (s), 157.4(s) Anal. calculated for C_21_H_16_N_4_O_2_S (MW 388.44): C, 64.93; H, 4.15; N, 14.42 %. Found: C, 65.22; H, 4.33; N, 14.70%.

### 5-Bromo-3-[6-(2,3-dihydro-benzo[1,4]dioxin-6-yl)-imidazo[2,1-b][1,3,4]thiadiazol-2-yl]-1H-indole (10c)

Yield: 55 %; light brown solid; m.p.: 327 °C; IR (cm^−1^): 3142 (NH);^1^H NMR (200 MHz, DMSO-*d_6_*) δ: 4.27 (4H, s, CH_2_), 6.89 (1H, d, *J* = 8.0 Hz, Ar), 7.32 (1H, s, Ar), 7.35 (1H, s, Ar), 7.42 (1H, d, *J* = 8.6 Hz, Ar), 7.51 (1H, d, *J* = 8.6 Hz,Ar), 8.32 (1H, s, Ar), 8.37 (1H, d, *J* = 1.3 Hz, Ar), 8.64 (1H, s, Ar), 12.21 (1H, s, NH). ^13^C NMR (50 MHz, DMSO-*d_6_*) δ: 64.6 (2xt), 106.7 (s), 110.3 (d), 113.6 (d), 114.5 (s),115.1 (d), 117.7 (d), 118.1 (d), 123.1 (d), 125.9 (s), 126.7 (d), 128.1 (s), 131.1 (d), 135.9 (s), 143.0 (s), 143.2 (s), 143.9 (s), 145.1 (s), 156.8 (s). Anal.calculated for C_20_H_13_BrN_4_O_2_S (MW 453.32): C, 52.99; H, 2.89; N, 12.36%. Found: C, 53.25; H, 2.97; N, 12.60%.

### 3-[6-(2,3-Dihydro-benzo[1,4]dioxin-6-yl)-imidazo[2,1-b][1,3,4]thiadiazol-2-yl]-5-fluoro-1-methyl-1H-indole (10d)

Yield: 58 %; yellow solid; m.p.: 257 °C; ^1^H NMR (200 MHz, DMSO-*d_6_*) δ: 3.88 (3H, s, CH_3_) 4.28 (4H, s, CH_2_), 6.88 (1H, d, *J* = 8.2 Hz, Ar), 7.33 (2H, d, *J* = 7.8 Hz, Ar), 7.47 (1H, d, *J* = 8.7 Hz, Ar), 7.59 (1H, d, *J* = 8.7 Hz, Ar), 8.31 (2H, d, *J* = 9.7 Hz, Ar), 8.58 (1H, s, Ar). ^13^C NMR (50 MHz, DMSO-*d_6_*) δ: 33.8 (q), 64.3 (t), 65.2 (t), 102.8 (s), 105.6 (d), 113.7 (d), 114.9 (d), 115.8 (d), 118.3 (d), 122.6 (d), 123.4 (d), 124.4 (s), 126.2 (s), 134.5 (s), 136.5 (d), 143.9 (s), 146.8 (s), 148.0 (s), 150.1 (s), 156.0 (s), 159.8 (s). Anal. calculated for C_21_H_15_BrN_4_O_2_S (MW 467.34): C, 53.9; H, 3.24, N, 11.99%. Found: C, 53.81; H, 3.35; N, 12.23%.

### 5-Chloro-3-[6-(2,3-dihydro-benzo[1,4]dioxin-6-yl)-imidazo[2,1-b][1,3,4]thiadiazol-2-yl]-1H-indole (10e)

Yield: 75 %; yellow solid; m.p.: 340 °C; IR (cm^−1^): 3136 (NH);^1^H NMR (200 MHz, DMSO-*d_6_*) δ: 4.28 (4H, s, CH_2_), 6.91 (1H, d, *J* = 7.5 Hz, Ar), 7.43 – 7.27 (3H, m, Ar), 7.57 (1H, d, *J* = 7.6 Hz, Ar), 8.17 (1H, s, Ar), 8.40 (1H, s, Ar), 8.65 (1H, s, Ar), 12.30 (1H, s, NH). ^13^C NMR (50 MHz, DMSO-*d_6_*) δ: 64.6 (2xt), 106.8 (s), 110.2 (d), 113.6 (d), 114.6 (d), 117.7 (d), 118.1 (d), 120.0 (d), 123.7 (d), 125.3 (s), 126.5 (s), 128.0 (s), 131.2 (d), 135.7 (s), 143.1 (s), 143.2 (s), 143.9 (s), 145.1 (s), 156.8 (s). Anal. calculated for C_20_H_13_ClN_4_O_2_S (MW 408.86): C, 58.75; H, 3.20; N, 13.70%. Found: C, 58.91; H, 3.48; N, 13.93%.

### 5-Chloro-3-[6-(2,3-dihydro-benzo[1,4]dioxin-6-yl)-imidazo[2,1-b][1,3,4]thiadiazol-2-yl]-1-methyl-1H-indole (10f)

Yield: 78%; brown solid; m.p.: 270 °C; ^1^H NMR (200 MHz, DMSO-*d_6_*) δ: 3.88 (3H, s, CH_3_), 4.27 (4H, s, CH_2_), 6.89 (1H, d, *J* = 8.1 Hz, Ar), 7.34 (3H, m, Ar), 7.64 (1H, d, *J* = 8.7 Hz, Ar), 8.13 (1H, s, Ar), 8.36 (1H, s, Ar), 8.59 (1H, s, Ar). ^13^C NMR (50 MHz, DMSO-*d_6_*) δ: 33.8 (q), 64.4 (2xt), 105.7 (d), 110.3 (d), 113.3 (d), 113.6 (d), 117.7 (s), 118.1 (d), 120.1 (d), 123.7 (s),125.4 (d), 126.9 (s), 127.9 (d), 134.6 (s), 136.3 (s), 142.9 (s), 143.2 (s), 143.9 (s), 145.1 (s), 156.4 (s). Anal. calculated for C_21_H_15_ClN_4_O_2_S (MW 422.89): C, 59.64; H, 3.58; N, 13.25%. Found: C, 59.81; H, 3.40; N, 13.50%.

### 3-[6-(2,3-Dihydro-benzo[1,4]dioxin-6-yl)-imidazo[2,1-b][1,3,4]thiadiazol-2-yl]-5-fluoro-1H-indole (10g)

Yield: 70 %; beige solid; m.p.: 330 °C; IR (cm^−1^): 3193 (NH);^1^H NMR (200 MHz, DMSO-*d_6_*) δ: 4.28 (4H, s, CH_2_), 6.90 (1H, d, *J* = 8.0 Hz, Ar), 7.16 (1H, t, *J* = 8.5 Hz, Ar), 7.35 (2H, d, *J* = 8.5 Hz, Ar), 7.56 (1H, dd, *J* = 8.5, 4.2 Hz, Ar), 7.84 (1H, d, *J* = 9.8 Hz, Ar), 8.37 (1H, s, Ar), 8.59 (1H, s, Ar), 12.21 (1H, s, NH).^13^C NMR (50 MHz, DMSO-*d_6_*)δ: 64.6 (2xt), 105.8 (d), 107.2 (s), 110.2 (d), 111.9 (d), 113.6 (d), 114.3 (d), 117.7 (d), 118.1 (d), 124.5 (s), 128.1 (s), 131.4 (d), 133.8 (s), 143.2 (s), 143.9 (s), 145.1 (s), 156.9 (s), 157.1 (s), 160.2 (s). Anal. calculated for C_20_H_13_FN_4_O_2_S (MW 392.41): C, 61.22; H, 3.34; N, 14.28%. Found: C, 61.48; H, 3.58; N, 14.07%.

### 3-[6-(2,3-Dihydro-benzo[1,4]dioxin-6-yl)-imidazo[2,1-b][1,3,4]thiadiazol-2-yl]-5-fluoro-1-methyl-1H-indole (10h)

Yield: 65%; beige solid; mp: 271 °C; ^1^H NMR (200 MHz, DMSO-*d_6_*) δ: 3.89 (3H, s, CH_3_) 4.28 (4H, s, CH_2_), 6.89 (1H, d, *J* = 8.0 Hz, Ar), 7.22 (1H, t, *J* = 8.5 Hz, Ar), 7.35 (2H, s, Ar), 7.63 (1H, dd, *J* = 8.1, 3.7 Hz, Ar), 7.82 (1H,d, *J* = 9.4 Hz, Ar), 8.35 (1H, s, Ar), 8.54 (1H, s, Ar). ^13^C NMR (50 MHz, DMSO-*d_6_*) δ: 33.9 (q), 64.6 (2xt), 105.6 (d), 105.9 (s), 106.1 (d), 110.3 (d), 111.8 (d), 112.0 (d), 113.1 (d), 114.5 (s), 124.9 (s), 134.5(d), 134.9 (s), 137.7(d), 142.4 (s), 143.6 (s), 149.7 (s), 157.0 (s), 157.4 (s),160.6 (s). Anal. calculated for C_21_H_15_FN_4_O_2_S (MW 406.43): C, 62.06; H, 3.72; N, 13.79%. Found: C, 62.32; H, 3.98; N, 13.85%.

### 3-[6-(2,3-Dihydro-benzo[1,4]dioxin-6-yl)-imidazo[2,1-b][1,3,4]thiadiazol-2-yl]-5-methoxy-1H-indole (10i)

Yield: 67 %; off-white solid; m.p.: 265 °C; IR (cm^−1^): 3197 (NH); ^1^H NMR (200 MHz, DMSO-*d_6_*) δ: 3.85 (3H, d, *J* = 2.6 Hz, CH_3_), 4.27 (4H, s, CH_2_), 6.98 – 6.86 (2H, m, Ar), 7.35 (2H, dd, *J* = 7.7, 2.5 Hz, Ar), 7.44 (1H, dd, *J* = 8.8, 2.5 Hz, Ar), 7.63 (1H, s, Ar), 8.23 (1H, s, Ar), 8.59 (1H, d, *J* = 2.7 Hz, Ar), 11.98 (1H, s, NH). ^13^C NMR (50 MHz, DMSO-*d_6_*) δ: 55.9 (q), 64.6 (2xt), 102.8 (d), 106.8 (s), 110.2 (d), 113.5 (d), 113.6 (d), 113.8 (d), 117.7 (d), 118.1 (d), 124.9(s), 128.0 (s), 130.0 (d), 132.1 (s), 143.0 (s), 143.2 (s), 143.9 (s), 144.9 (s), 155.6 (s), 157.5 (s). Anal. calculated for C_21_H_16_N_4_O_3_S (MW 404.44): C, 62.36; H, 3.99; N, 13.85%. Found: C, 62.55; H, 3.71; N, 13.98%.

### 3-[6-(2,3-Dihydro-benzo[1,4]dioxin-6-yl)-imidazo[2,1-b][1,3,4]thiadiazol-2-yl]-5-methoxy-1-methyl-1H-indole (10j)

Yield: 47 %; light brown solid; m.p.: 238 °C; ^1^H NMR (200 MHz, DMSO-d6) δ: 3.86 (6H, s, CH_3_), 4.27 (4H, s, CH_2_), 6.88 (1H, d, *J* = 8.3 Hz, Ar), 6.99 (1H, dd, J = 8.9, 2.2 Hz, Ar), 7.35 (2H, d, *J* = 7.9 Hz, Ar), 7.51 (1H, d, *J* = 8.9 Hz, Ar), 7.62 (1H, d, *J* = 2.1 Hz, Ar), 8.22 (1H, s, Ar), 8.55 (1H, s, Ar). ^13^C NMR (50 MHz, DMSO-d6) δ: 33.7 (q), 55.9 (q), 64.6 (2xt), 102.9 (d), 105.6 (s), 110.2 (d), 112.3 (d), 113.4 (2xd), 113.6 (d), 117.7 (d), 118.1 (d), 125.2 (s), 128.1 (s), 132.9 (s), 133.4 (s), 143.1 (s), 143.9 (s), 145.0 (s), 155.9 (s), 156.9 (s). Anal. calculated for C_22_H_18_N_4_O_3_S (MW 418.47): C, 63.14; H, 4.34; N, 13.39%. Found: C, 63.31; H, 4.62; N, 13.53%.

### 3-(6-Furan-2-yl-imidazo[2,1-b][1,3,4]thiadiazol-2-yl)-1H-indole (10k)

Yield: 47 %; white solid; m.p.: 218°C; IR (cm^−1^) 3152 (NH); ^1^H NMR (200 MHz, DMSO-*d_6_*) δ: 6.59 (1H, d, *J* = 1.3 Hz, Ar), 6.71 (1H, d, *J* = 3.4 Hz, Ar), 7.34 – 7.25 (2H, m, Ar), 7.55 (1H, dd, *J* = 6.8, 2.7 Hz, Ar), 7.70 (1H, d, *J* = 2.1 Hz, Ar), 8.18 (1H, dd, *J* = 4.9, 2.6 Hz, Ar), 8.33 (1H, d, *J* = 2.8 Hz, Ar), 8.41 (1H, d, *J* = 4.3 Hz, Ar), 12.14 (1H, s, NH). ^13^C NMR (50 MHz, DMSO-*d_6_*) δ: 105.6 (d), 106.9 (s), 110.22(d), 112.1 (d), 113.0 (d), 120.9 (d), 121.9 (d), 123.7 (d), 124.2 (s), 129.9 (d), 137.2 (s), 137.6 (s), 142.4 (d), 143.8 (s), 149.7 (s), 157.8 (s). Anal. calculated for C_16_H_10_N_4_OS (MW: 306.34): C, 62.73; H, 3.29; N, 18.29%. Found: C, 62.98; H, 3.43; N, 18.51%.

### 3-(6-Furan-2-yl-imidazo[2,1-b][1,3,4]thiadiazol-2-yl)-1-methyl-1H-indole (10l)

Yield: 56 %; yellow solid; m.p.: 188 °C; ^1^H NMR (200 MHz, DMSO-*d_6_*) δ: 3.90 (3H, s, CH_3_), 6.59 (1H, s, Ar), 6.70 (1H, s, Ar), 7.44 – 7.26 (2H, m, Ar), 7.62 (1H, d, *J* = 7.3 Hz, Ar), 7.71 (1H, s, Ar), 8.18 (1H, d, *J* = 6.3 Hz, Ar), 8.34 (1H, s, Ar), 8.40 (1H, s, Ar). ^13^C NMR (50 MHz, DMSO-*d_6_*) δ: 33.6 (q), 105.6 (d), 105.9 (s), 110.2 (d), 111.5 (d), 112.1 (d), 121.0 (d), 122.3 (d), 123.7 (d), 124.5 (s), 133.5 (d), 137.6 (s), 137.7 (s), 142.4 (d), 143.7 (s), 149.7 (s), 157.4 (s). Anal. calculated for C_17_H_12_N_4_OS (MW: 320.07): C, 63.73; H, 3.78; N, 17.49%. Found: C, 63.91; H, 3.58; N, 17.67%.

### 5-Bromo-3-(6-furan-2-yl-imidazo[2,1-b][1,3,4]thiadiazol-2-yl)-1H-indole (10m)

Yield: 55 %; brown solid; m.p.: 285 °C; IR (cm^−1^) 3147 (NH); ^1^H NMR (200 MHz, DMSO-*d_6_*) δ: 6.59 (1H, d, *J* = 1.4 Hz, Ar), 6.69 (1H, d, *J* = 3.0 Hz, Ar), 7.42 (1H, d, *J* = 7.6 Hz, Ar), 7.52 (1H, dd, *J* = 8.6, 3.2 Hz, Ar), 7.70 (1H, s, Ar), 8.32 (1H, s, Ar), 8.39 (1H, d, *J* = 3.0 Hz, Ar), 8.48 (1H, d, *J* = 3.6 Hz, Ar), 12.31 (1H, s, NH). ^13^C NMR (50 MHz, DMSO-*d_6_*) δ: 105.6 (d), 106.6 (s), 110.4 (d), 112.1 (d), 114.6 (s), 115.1 (d), 123.1 (d), 125.9 (s), 126.3 (d), 131.3 (d), 135.9 (s), 137.7 (s), 142.4 (d), 143.7 (s), 149.7 (s), 157.3 (s). Anal. calculated for C_16_H_9_BrN_4_OS (MW: 385.24): C, 49.88; H, 2.35; N, 14.54%. Found: C, 49.95; H, 2.61; N, 14.33%.

### 5-Bromo-3-(6-furan-2-yl-imidazo[2,1-b][1,3,4]thiadiazol-2-yl)-1-methyl-1H-indole (10n)

Yield: 48 %; pale yellow solid; m.p.: 245 °C; ^1^H NMR (200 MHz, DMSO-*d_6_*) δ: 3.89 (3H, s, CH_3_), 6.58 (1H, dd, *J* = 3.0, 1.6 Hz, Ar), 6.69 (1H, d, *J* = 3.1 Hz, Ar), 7.48 (1H, dd, *J* = 8.7, 1.5 Hz, Ar), 7.60 (1H, d, *J* = 8.8 Hz, Ar) 7.69 (1H, s, Ar), 8.30 (1H, s, Ar), 8.37 (1H, s, Ar), 8.44 (1H, s, Ar). ^13^C NMR (50 MHz, DMSO-*d_6_*) δ: 33.8 (q), 105.5 (d), 105.7 (s), 110.4 (d), 112.0 (d), 113.7 (s), 115.0 (d), 123.1 (d), 126.0 (s), 126.3 (d), 134.7 (d), 136.5 (d), 137.6 (s), 142.4 (s), 143.5 (s), 149.7 (s), 156.9 (s). Anal. calculated for C_17_H_11_BrN_4_OS (MW: 399.26): C, 51.14; H, 2.78; N, 14.03%. Found: C, 51.45; H, 2.92; N, 14.15%.

### 5-Chloro-3-(6-furan-2-yl-imidazo[2,1-b][1,3,4]thiadiazol-2-yl)-1H-indole (10o)

Yield: 52 %; yellow solid; m.p.: 248°C; IR (cm^−1^) 3150 (NH); ^1^H NMR (200 MHz, DMSO-*d_6_*) δ: 6.58 (1H, dd, *J* = 3.2, 1.7 Hz, Ar), 6.69 (1H, d, *J* = 3.1 Hz, Ar), 7.31 (1H, dd, J = 8.6, 2.0 Hz, Ar), 7.57 (1H, d, *J* = 8.7 Hz, Ar), 7.70 (1H, s, Ar), 8.17 (1H, d, *J* = 1.7 Hz, Ar), 8.40 (1H, d, *J* = 3.0 Hz, Ar), 8.46 (1H, d, *J* = 3.5 Hz, Ar), 12.31 (1H, s, NH). ^13^C NMR (50 MHz, DMSO-*d_6_*) δ: 105.7 (d), 106.8 (s), 107.8 (d), 110.4 (d), 114.6 (s), 114.7 (d), 120.1 (d), 123.8 (d), 125.3 (s), 126.3 (s), 126.6 (s), 131.5 (d), 135.7 (s), 137.7 (s),143.7 (d), 149.7 (s). Anal. calculated for C_16_H_9_ClN_4_OS (MW: 340.79): C, 56.39; H, 2.66; N, 16.44%. Found: C, 56.51; H, 2.86; N, 16.63%.

### 5-Chloro-3-(6-furan-2-yl-imidazo[2,1-b][1,3,4]thiadiazol-2-yl)-1-methyl-1H-indole (10p)

Yield: 55 %; white solid; m.p.: 253 °C; 1H NMR (200 MHz, DMSO-*d_6_*) δ: 3.90 (3H, s, CH_3_), 6.58 (1H, dd, *J* = 3.0, 1.7 Hz, Ar), 6.69 (1H, d, *J* = 3.0 Hz, Ar), 7.49 (1H, dd, *J* = 8.7, 1.5 Hz, Ar),7.65 – 7.56 (1H, m, Ar), 7.68 (1H, d, *J* = 3.4 Hz, Ar), 8.31 (1H, s, Ar), 8.37 (1H, d, *J* = 4.9 Hz, Ar), 8.42 (1H, d, *J* = 3.5 Hz, Ar). ^13^C NMR (50 MHz, DMSO-*d_6_*) δ: 33.8 (q), 105.5 (d), 105.7 (s), 110.4 (d), 112.0 (d), 113.3 (s), 115.0 (d), 120.1 (d), 126.0 (s), 127.0 (d), 134.7 (d), 136.3 (d), 137.7 (s), 142.4 (s), 143.6 (s), 149.7 (s), 156.8 (s). Anal. Calculated for C_17_H_11_ClN_4_OS (MW: 354.81): C, 57.55; H, 3.12; N, 15.79%. Found: C, 57.84; H, 3.31; N, 15.91%.

### 5-Fluoro-3-(6-furan-2-yl-imidazo[2,1-b][1,3,4]thiadiazol-2-yl)-1H-indole (10q)

Yield: 50 %; off-white solid; m.p.: 265 °C; IR (cm^−1^) 3153 (NH);^1^H NMR (200 MHz, DMSO-*d_6_*) δ: 6.58 (1H, d, *J* = 1.3 Hz, Ar), 6.69 (1H, d, *J* = 2.9 Hz, Ar), 7.17 – 7.11 (1H, m, Ar), 7.58 – 7.53 (1H, m, Ar), 7.70 (1H, s, Ar), 7.86 (1H, dd, *J* = 9.5, 1.7 Hz, Ar), 8.39 (1H, d, *J* = 2.8 Hz, Ar), 8.41 (1H, s, Ar), 12.22 (1H, s, NH). ^13^C NMR (50 MHz, DMSO-*d_6_*) δ: 105.6 (d), 105.9 (d), 107.1 (s), 110.3 (d), 111.8 (d), 112.1 (d), 114.4 (d), 124.5 (s), 124.7 (s), 131.6 (d), 133.8 (s), 137.6 (s), 142.4 (d), 149.7 (s), 157.0 (s), 157.4 (s), 160.3 (s), 166.9 (s). Anal. Calculated for C_16_H_9_FN_4_OS (MW:324.33): C, 59.25; H, 2.80; N, 17.27%. Found: C, 59.43; H, 2.93; N, 17.41%.

### 5-Fluoro-3-(6-furan-2-yl-imidazo[2,1-b][1,3,4]thiadiazol-2-yl)-1-methyl-1H-indole (10r)

Yield: 65%; off-white solid; m.p.:246 °C; ^1^H NMR (200 MHz, DMSO-*d_6_*) δ: 3.90 (3H, s, CH_3_), 6.58 (1H, s, Ar), 6.69 (1H, d, *J* = 3.0 Hz, Ar), 7.22 (1H, td, *J* = 9.1, 2.0 Hz, Ar), 7.64 (1H, dd, *J* = 9.0, 4.4 Hz, Ar), 7.69 (1H, s, Ar), 7.85 (1H, dd, *J* = 9.7, 2.0 Hz, Ar), 8.37 (2H, s, Ar). ^13^C NMR (50 MHz, DMSO-*d_6_*) δ: 33.9 (q), 105.6 (d), 105.8 (s), 106.1 (d), 110.3 (d), 111.8 (d),112.1 (d), 113.0 (d), 124.7(s), 134.5 (s), 134.9 (d), 137.7 (s), 142.4 (d), 149.7 (s), 157.0 (s), 157.4 (s), 160.5 (s). Anal. Calculated for C_17_H_11_FN_4_OS (MW: 338.36): C, 60.34; H, 3.28; N, 16.56%. Found: C, 60.52; H, 3.35; N, 16.72%.

### 3-(6-Furan-2-yl-imidazo[2,1-b][1,3,4]thiadiazol-2-yl)-5-methoxy-1H-indole(10s)

Yield: 50 %; yellow solid; m.p.: 252 °C; IR (cm^−1^) 3147 (NH); ^1^H NMR (200 MHz, DMSO-*d_6_*) δ: 3.85 (3H, s, CH_3_), 6.57-6.59 (1H, m, Ar), 6.69 (1H, d, *J* = 3.2 Hz, Ar), 6.94 (1H, dd, *J* = 8.8, 2.4 Hz, Ar), 7.44 (1,H, d, *J* = 8.8 Hz, Ar), 7.64 (1H, d, *J* = 2.3 Hz, Ar), 7.69 (1H, s, Ar), 8.24 (1H, d, *J* = 3.0 Hz, Ar), 8.40 (1H, s, Ar), 11.99 (1H, s, NH). ^13^C NMR (50 MHz, DMSO-*d_6_*) δ: 55.9 (q), 103.0 (d), 105.6 (d), 106.8 (s), 110.3 (d), 112.0 (d), 113.5 (d), 113.8 (d), 124.9 (s), 130.2 (d), 132.2 (s), 137.6 (s), 142.3 (d), 143.7 (s), 149.8 (s), 155.7 (s), 157.9 (s). Anal. calculated for C_17_H_12_N_4_O_2_S (MW: 336.37): C, 60.70; H, 3.60; N, 16.66%. Found: C, 60.95; H, 3.79; N, 16.83%.

### 3-(6-Furan-2-yl-imidazo[2,1-b][1,3,4]thiadiazol-2-yl)-5-methoxy-1-methyl-1H-indole (10t)

Yield: 54 % ; brown solid; m.p.: 167 °C; ^1^H NMR (200 MHz, DMSO-*d_6_*) δ: 3.86 (6H, s, CH_3_), 6.59 (1H, s, Ar), 6.69 (1H, s, Ar), 7.00 (1H, d, *J* = 8.8 Hz, Ar), 7.53 (1H, d, *J* = 9.0 Hz, Ar), 7.64 (1H, s, Ar), 7.70 (1H, s, Ar), 8.26 (1H, s, Ar), 8.39 (1H, s, Ar). ^13^C NMR (50 MHz, DMSO-*d_6_*) δ: 33.7 (q), 56.0 (q), 102.1 (s), 103.2 (d), 105.6 (d), 110.3 (d), 112.0 (d), 112.4 (d), 113.3 (d), 125.2 (s), 132.9 (s), 133.6 (d), 137.5 (s), 141.6 (s), 142.4 (d), 149.7 (s), 155.9 (s), 157.5 (s). Anal. calculated for C_18_H_14_N_4_O_2_S (MW: 350.39): C, 61.70; H, 4.03; N, 15.99%. Found: C, 61.93; H, 4.21; N, 15.74%.

### Biology

#### Drugs and chemicals

For the biological assays the newly synthesized compounds **10** were dissolved in DMSO. The medium, foetal bovine serum (FBS), penicillin (50 IU mL^−1^) and streptomycin (50 mg mL^−1^) were purchased from Gibco (Gaithersburg, MD, USA), whereas the other chemicals were acquired from Sigma (Zwijndrecht, the Netherlands).

#### Cell culture

The cells were cultured in RPMI-1640 (Roswell Park Memorial Institute 1640) enriched with 10% heat-inactivated newborn calf serum (NBCS), 1% penicillin/streptomycin, or in DMEM (Dulbecco’s Modified Eagle’s Medium), added with 10% heat-inactivated NBCS. They were maintained in a humidified environment containing 5% CO2 and 95% air at 37 °C and harvested using trypsin-EDTA.

#### Cell growth inhibition

The cytotoxic activity of the imidazothiadiazole derivatives **10** against differentiated pancreatic cancer cells was assessed using the sulforhodamine B (SRB) chemosensitivity assay. Triplicate wells of 96-well flat-bottom plates were seeded with cells in volumes of 100 mL, with different cell densities for each cell line (3 × 10^3^ cells/well for SUIT-2, PANC-1, PANC-1 GR, PATU-T, and PDAC-3; 4 × 10^3^ cells/well for BxPC-3; 5 × 10^3^ cells/well for CAPAN-1; and 8 × 10^3^ cells/well for the normal fibroblasts Hs27, used to evaluate the selectivity index (SI)). After incubating for 24 hours at 37 °C, cells were treated with various concentrations of the compounds (100 µL) for 72 hours at 37 °C, 5% CO_2_, and 95% humidity.

Upon completion of the incubation period, the cells were treated with 25 µL of 50% cold trichloroacetic acid for fixation and maintained at 4 °C for a minimum of 60 minutes. Subsequently, the plates were emptied, gently rinsed with deionized water, dried overnight at room temperature (RT), and stained with 50 µL of 0.4% SRB solution in 1% acetic acid for 15 minutes at RT. After removing excess SRB stain, the plates were washed with a 1% acetic acid solution and allowed to air-dry overnight at RT. The SRB staining was dissolved in 150 µL of tris(hydroxymethyl)aminomethane solution pH = 8.8 (TRIS base), and the absorbance, at wavelengths of 490 nm and 540 nm, was evaluated. Cell growth inhibition was determined as the percentage of drug treated cells compared to vehicle-treated cells (“untreated cells or control”) OD (adjusted for OD before drug exposure, “day-0”).

The percentage of cell growth was calculated by comparing the average optical density of the growth in control wells and of the sample wells, through the following equation:

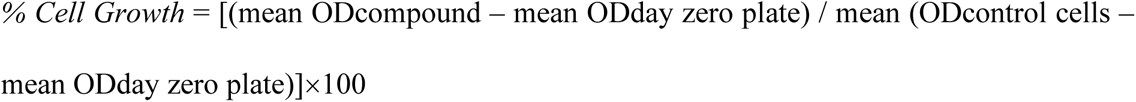

The outcomes achieved were calibrated using the day zero plate (wells harboring cells grown for just 24 hours) and standardized by the control cells (wells containing untreated cells) to obtain the viable cells rate. The 50% inhibitory concentration of cell growth (IC_50_) was computed through non-linear least squares curve fitting (GraphPad PRISM version 9, Intuitive Software for Science, San Diego, CA). In the NCI procedure, the IC_50_ is referred to as the GI_50_, signifying the concentration at which 50% growth inhibition is attained. Data was expressed as mean values ± SEM.

Initially, the cells were exposed to three different screening concentrations (0.1, 1and 10 μM) of each compound **10** for a period of 72 hours. Subsequently, the SRB assay was executed to ascertain the inhibition of cell proliferation induced by the most active compunds. To determine the IC_50_ values for each PDAC cell model, cells were treated with eight escalating concentrations (ranging from 0.3 μM to 40 μM) of compounds **10f**, **10k**, **10l**, and **10q** for 72 hours.

#### Wound-healing assay

The *in vitro* scratch wound-healing assay was conducted following a previously established method [34]. BxPC-3, PATU-T, SUIT-2, CAPAN-1, PANC-1, PANC-1 GR, and PDAC-3 cells were seeded in 96-well flat-bottom plates at a density of 3-5 × 10^4^ cells/well in a volume of 100 µL. After 24 hours of pre-incubation at 37 °C, 5% CO_2_, and 95% humidity, scratches of consistent width were created on the cell monolayers using a specialized multi-needle tool. Following the removal of detached cells by washing with phosphate-buffered saline, the control wells received only medium, while the experimental wells were supplemented with the compounds of interest. The wound confluence was monitored at specific time points (T = 0, 4, 8, 20, and 24 hours after treatment) using phase-contrast microscopy integrated with the Leica DMI300B migration station and Universal Grab 6.3 software from Digital Cell Imaging Labs (Keerbergen, Belgium). Images were captured, and the wound closure was analyzed using the Scratch Assay 6.2 software from Digital Cell Imaging Labs.

#### PamChip® kinase activity profiling

A PamChip array comprising 144 kinase peptide substrates was employed to assess the alteration in tyrosine kinase activity upon the application of **10l** compound. This study was conducted using SUIT-2 and PDAC-3 cells in biological triplicates, involving three untreated samples and three samples treated with **10l**, as outlined previously [35].

The cells were cultured in 25 cm^2^ flasks until they attained 80% confluence, maintaining a temperature of 37 °C and a CO_2_ concentration of 5%. Following this, both cell lines were exposed to 5 times the IC_50_ concentration of **10l**, while the control cells underwent medium replacement with fresh medium. The treatment duration was 2 hours, after which the cells were detached, and 10^6^ cells/mL were harvasted from each replicate in 1.5 mL tubes. To lyse the cells, 100 µL of M-PER lysis buffer per 10^6^ cells was utilized, comprising M-PER Mammalian Extraction Buffer (Thermo Scientific, Rockford, IL, USA), Halt protease inhibitor cocktail (Complete Mini EDTA-free Protease Inhibitor Cocktail, Roche #11836170001) diluted 1:100, and Halt Phosphatase Inhibitor Cocktail (Thermo Fisher #78420) also diluted 1:100. The lysis process took at least 15 minutes at 4 °C. The lysates were then collected in 1.5 mL tubes and centrifuged at 16,000g for 15 minutes at 4 °C. The resulting supernatants were gathered and stored at −80 °C until further use. The protein concentration of the samples was estimated employing the Bio-Rad protein assay, which is based on the Bradford method (BioRad, Hercules, CA). The PamChips were prepared in order to reach a concentration of 10 µg protein/array. They were then added to the MasterMix (PamGene reagent kit 32116) which included PK buffer 10x, BSA solution 100x, PTK additive 10x, 1 M DTT solution, Complete Mini EDTA-free Protease Inhibitor Cocktail, Halt Phosphatase Inhibitor Cocktail 400x (Thermo Fisher), PY-20-FITC (fluorescent labeled antibody), and 4 mM ATP solution. The samples were added to the MasterMix promptly before loading onto the chip. Prior to sample loading, the PamStation®12 instrument performed a blocking step using 25 µL of 2% BSA on each array, followed by three wash steps with PK buffer. Subsequently, 40 µL of each sample mixture was loaded in duplicate onto the arrays. Throughout the 30 °C incubation, the sample mixture underwent pumping up and down through the array once every minute for a total of 60 cycles. Fluorescent imaging of each array was repeated using a 12-bit CCD camera, which monitored fluorescence intensities in real time.

The intensity of each spot at the end point was evaluated through an open-source software (ScanAnalyze) and then nomralized to account for local background noise. As the negative control yielded a negative value, all intensities were shifted minimally to ensure the negative control value was set to zero. For the differential analysis, a Student t-test was conducted in R (version 3.6.1), and the resulting p-values were corrected for false discovery rate (FDR). Peptides reaching a cutoff of FDR < 0.01 were considered statistically significant. Visualization of differentially phosphorylated protein was performed in Cytoscape (version 3.5.0).

#### FAK ELISA assay

The quantification of pFAK (phosphorylated focal adhesion kinase) at tyrosine residue 397 was carried out using an Enzyme-Linked Immunosorbent Assay (ELISA). The specific ELISA kit employed was the InvitrogenTM FAK[pY397] ELISA Kit (Cat. # KHO0441), following the manufacturer’s instructions. Supernatants from SUIT-2and PDAC-3 cells were collected 24 hours after treatment with imidazothiadiazole compounds **10k and 10l** at a concentration corresponding to 5 times the IC_50_ values. The absorbance of the samples was measured at 450 nm. Additionally, we conducted a parallel ELISA test using the well-known FAK inhibitor defactinibat 5 times IC_50_, which effectively inhibited FAK phosphorylation, as described previously [36].

#### Cell cycle assay

Cell cycle stage was analyzed by flow cytometry. Cells were seeded in 6 wells plate (2.5 × 10^5^ cells/well). After an overnight incubation at 37 °C, the cells were treated with the compound **10l** at 5 times IC_50_, and incubated for 24 hours. After treatment, the cells were harvested by trypsinization (0.3 mL/well of trypsine-EDTA), incubated until the cells were detached; cells were resuspended with 1.7 mL of medium and collected into FACS tubes. The samples were then centrifuged in order to get cells pellet (5 min at 1200 rpm). Finally, these pellets were fixed in ice-cold 70% ethanol, washed twice with phosphate-buffered saline (20 mM sodium phosphate pH 7.4, 150 mM NaCl) and incubated for 30 min at 37 °C with 50 µl of RNase (100 µg/mL) followed by incubation with 200 µl of propidium iodide solution (PI, 50 µg/mL). The cycle analysis was performed on the FACS (Fluorescence Activated Cell Sorting) Calibur instrument, to evaluate the effects on the cell cycle distribution and cell viability.

#### Evaluation of apoptosis

The evaluation of caspase-3 activity was chosen as a reliable method to measure apoptosis induction due to its well-established role as a key executioner caspase in the apoptotic pathway. To this goal we used a specific spectrofluorimetric activity assay (Human Active Caspase-3 Immunoassay Quantikine ELISA, Catalog Number KM300, R&D Systems, Inc., Minneapolis, MN). In summary, cells were seeded in 6-well plates at a density of 2 × 10^5^ cells/mL and treated with **10l** or defactinib at 5 × IC_50_ for 24 hours. Following drug exposure, cell extracts were prepared, diluted, and combined with the provided reagents following the manufacturers’ instructions. The absorbance was measured at 450 nm with a background subtraction at 540 nm. The relative caspase activity was determined by comparing the measurements to a standard curve generated using human recombinant caspase-3, as described previously [37].

#### Spheroids

PDAC-3 spheroids were grown in VWR®96-wells F-bottom plates (VWR, Cat No. 7342327) coated with 1.5% w/v agarose. Cells were seeded at the density of 2×10^4^ cells per well and the plate was centrifuged at 1200 rpm, for 5 min at RT to facilitate cell aggregation, finally cells were incubated at 37°C, 5% CO_2_ for 72 h in order to allow spheroids formation.

After 72 hours, before the treatment a picture of the plate was taken with an automated phase-contrast microscope DMI300B (Leica Microsystems, Eindhoven, Netherlands), and the subsequent pictures were taken every two days. Prior to treatment, automated phase-contrast microscope was used to capture an initial picture of the plate. Subsequent pictures were taken every two or three days.

After 72-hour incubation, the culture medium was replaced with drug-free medium for the control, whereas each testing group was treated with compounds of interest (at least 8 wells per condition). Compound **10l** and defactinib were used at 5 times the IC_50_,

Gemcitabine was used as an additional “positive control” at 150 nM, as previously reported [12]. The treatment and change of medium was repeated after four days, to ensure the availability of nutrients.

Pictures were analysed with ImageJ Software (U.S. National Institute of Health, Bethesda, Maryland, USA) to determine the area of the spheroids treated and compare it to the area of the untreated spheroids, as described previously [38].

#### Confocal experiment of spheroids

PDAC-3 spheroids were obtained as previously described. Spheroids were exposed to 150 nM gemcitabine, 3 and 15µM of **10l** after 3 days from seeding, and the treatment was refreshed every 3 days. Untreated spheroids served as controls. Viability and cell death were measured by live fluorescence confocal imaging. After the first treatment, spheroids were stained with a staining solution containing PBS 1X, FBS 1X, 1 µM Calcein AM, 0.32 µM Hoechst33342 and 3 µM of propidium iodide (PI) for 3 hours. Staining was repeated on day 8 (from seeding). Washing steps were avoided to preserve the integrity of the spheroids. For confocal imaging, z-stacks were acquired with a Nikon Eclipse Ti2 confocal microscope with a temperature and CO_2_ controlled incubator, using a 10x Plan dry objective. 4 images for each stack (∼11 stacks, steps of 20 µm) were stitched with NIS Elements 5.02.02 software. At least 3 spheroids per condition were imaged, on the day of the first treatment, and every 2 (on day 3 and 5) days in the first week and every 3 days in the second week (day8 and 11). Composite images were obtained by doing a z-projection with ImageJ. Scale bar is 400 µm.

#### *In vivo* experiment using subcutaneous mice

*In vivo* experiments were performed on mice purchased from Charles River Laboratories (Calco, Italy). The working protocol was approved by the local committees on animal experimentation of the VU University Medical Center (VUmc, Amsterdam, The Netherlands) and of the University of Pisa (Pisa, Italy), according to the 2010/63/EU European Community Council Directive for laboratory animal care.

*In vivo* experiments were performed on subcutaneous primary PDAC models. Subcutaneous tumors were obtained by cell suspension injection of 1 × 10^6^ primary PDAC-3 cells [39]. Mice were treated with compound **10l**, solubilised in Polyethylene glycol 400 (PEG400, Sigma-Aldrich, St. Louis, MO), at 25 mg/kg, 3 days/week for 2 weeks (formulation concentration: 25 mg/mL in PEG400, 100 µL of i.p. injection for a 25g mouse). When tumor volume reached an average size of 100 mm^3^ (day 5), the animals were randomly distributed into four groups (*n* = 5 tumours per treatment group) as follows: (1) untreated mice, (2) mice treated with gemcitabine alone at 100 mg/kg, 2 days (days 1 and 4) for 2 weeks (formulation concentration: 25 mg/mL in PBS, 100 µL of i.p. injection for a 25g mouse), (3) mice treated with **10l** solubilised in PEG400, at 25 mg/kg, 3 days (days 1, 3 and 5) for 2 weeks (formulation concentration: 12.5 mg/mL in PEG400). The concentration of **10l** was selected on the basis of previous experiments.

Tumor xenografts were measured as described previously [40]. Lidocaine was used as local anaesthetic on the skin of the mice. Mice were sacrificed before primary tumours and metastases can cause severe symptoms, via cervical dislocation.

#### Statistics

The *in vitro* experiments were performed in triplicates and repeated at least twice. The results reported in the figures are expressed as mean values ± standard error of the mean (SEM). Statistical analyses were carried out by two-way ANOVA followed by Bonferroni’s post-test (to adjust for multiple comparisons), using GraphPad-Prism version 9 (Intuitive Software for Science, San Diego, CA). All analyses were two-sided, and statistical significance was set at P < 0.05. *In vivo* statistical analysis was conducted with ANOVA analysis with Bonferroni correction for post hoc analysis to evaluate differences in tumor size in the different groups of animals. P < 0.05 was considered significant. The results were expressed as mean values ± standard deviation (SD).

## Results and Discussion

### Antiproliferative activity

The newly synthesized imidazo [2,1-*b*][1,3,4]thiadiazoles **10a-t** were tested by the NCI (Bethesda, MD) for the evaluation of their antitumor activity. They were initially prescreened, according to the NCI protocol, at one-dose (10 µM) on the full panel of 60 human cancer cell lines, derived from 9 cancer cell types and grouped into disease subpanels, including leukemia, non-small cell lung, colon, central nervous system, melanoma, ovarian, renal, prostate, and breast cancers.

Among the tested compounds, the furan analogues demonstrated superior potency compared to the corresponding benzodioxane derivatives. In particular compounds **10k**, **10l**, **10n**, and **10r**, meeting the criteria set by the NCI, underwent further screening across the full panel at five concentrations, ranging from 10^−4^ to 10^−8^ M, in order to determine their half maximal inhibitory concentrations (GI_50_). All four compounds exhibited significant antiproliferative activity against the tested cell lines, with GI_50_ values in the nanomolar or low micromolar range (Table 2). Compound **10l** displayed the highest potency, eliciting nanomolar GI_50_ values against fifteen different cancer cells. Notably, it exhibited GI_50_ values in the range from 0.45 to 3.09 µM and from 0.52 to 3.64 µM against the leukemia and colon cancer subpanels, respectively.

**Table 2.** Antiproliferative activity^a^ of compounds 10k, 10l, 10n and 10r expressed as GI_50_^b^ and TGI.

| Panel/Cell line | 10k |  | 10l |  | 10n |  | 10r |  |
| --- | --- | --- | --- | --- | --- | --- | --- | --- |
| | GI<sub>50</sub><br>(μM) | TGI<br>(μM) | GI<sub>50</sub><br>(μM) | TGI<br>(μM) | GI<sub>50</sub><br>(μM) | TGI<br>(μM) | GI<sub>50</sub><br>(μM) | TGI<br>(μM) |
| **Leukemia** |  |  |  |  |  |  |  |  |
| CCRF-CEM | 2.96 | 29.6 | 3.01 | >100 | 33.5 | >100 | 4.19 | 36.2 |
| HL-60(TB) | 2.10 | 89.8 | 0.92 | >100 | >100 | >100 | 2.81 | 10.0 |
| K-562 | 0.56 | >100 | 0.45 | >100 | 17.6 | >100 | 2.18 | 17.8 |
| MOLT-4 | 3.02 | >100 | 1.90 | >100 | 21.4 | 88.1 | 3.38 | 13.5 |
| RPMI-8226 | 3.19 | >100 | 3.09 | >100 | 4.66 | 83.0 | 4.06 | 73.7 |
| SR | 0.69 | >100 | 0.45 | >100 | 35.6 | >100 | 2.85 | 16.6 |
| **Non-Small Cell Lung Cancer** |  |  |  |  |  |  |  |  |
| A549/ATCC | 4.09 | >100 | 3.56 | >100 | 58.9 | 40.3 | 7.75 | 45.2 |
| EKVX | 1.76 | >100 | 1.56 | >100 | 10.6 | 32.9 | 5.15 | 40.2 |
| HOP-62 | 4.62 | >100 | 3.19 | >100 | 8.06 | 49.3 | 7.91 | 41.3 |
| HOP-92 | 6.39 | 63.2 | 14.7 | 98.7 | 2.09 | 10.6 | 4.45 | 22.7 |
| NCI-H226 | 24.0 | >100 | 22.3 | >100 | n.t. | n.t. | n.t. | n.t. |
| NCI-H23 | 2.82 | 39.8 | 3.24 | >100 | 7.83 | 27.1 | 7.45 | 39.4 |
| NCI-H322M | 3.46 | >100 | 6.45 | >100 | 6.97 | 52.9 | 10.9 | 49.7 |
| NCI-H460 | 3.71 | >100 | 2.42 | >100 | 7.04 | 22.7 | 3.72 | 16.1 |
| NCI-H522 | 1.97 | 18.2 | 0.72 | 17.7 | 17.0 | 48.1 | 5.17 | 35.2 |
| **Colon Cancer** |  |  |  |  |  |  |  |  |
| COLO 205 | 2.36 | 11.2 | 1.22 | 6.28 | 39.0 | >100 | 6.67 | 39.2 |
| HCC-2998 | 4.14 | 41.2 | 3.64 | 25.8 | 17.5 | 78.3 | 5.41 | 19.2 |
| HCT-116 | 2.42 | 27.8 | 0.71 | >100 | 2.74 | 8.65 | 4.39 | 16.2 |
| HCT-15 | 1.55 | >100 | 0.52 | 37.2 | 4.09 | 24.1 | 3.20 | 16.7 |
| HT29 | 2.79 | 69.6 | 0.67 | >100 | 15.2 | 64.1 | 4.44 | 20.2 |
| KM12 | 3.07 | 25.7 | 1.71 | >100 | 12.7 | 48.1 | 2.99 | 10.3 |
| SW-620 | 2.46 | >100 | 0.72 | >100 | 17.2 | 41.3 | 4.31 | 49.3 |
| **CNS Cancer** |  |  |  |  |  |  |  |  |
| SF-268 | 9.10 | >100 | 22.5 | >100 | 6.76 | 30.3 | 10.3 | 42.8 |
| SF-295 | 3.00 | 27.1 | 0.92 | 15.4 | 16.8 | 58.4 | 6.14 | 21.1 |
| SF-539 | 2.82 | 12.9 | 1.66 | 4.98 | 4.72 | 20.4 | 5.92 | 19.9 |
| SNB-19 | 6.68 | >100 | 4.88 | >100 | 12.6 | 27.1 | 11.0 | 30.4 |
| SNB-75 | 1.42 | 8.39 | 0.48 | 6.29 | 21.1 | 61.5 | 8.45 | 34.5 |
| U251 | 3.64 | >100 | 3.69 | >100 | 5.61 | 66.1 | 6.52 | 29.8 |
| **Melanoma** |  |  |  |  |  |  |  |  |
| LOX IMVI | 2.98 | 33.5 | 2.40 | 28.7 | 3.65 | 14.4 | 4.88 | 17.0 |
| MALME-3M | 2.41 | >100 | 1.12 | 27.0 | 7.64 | 29.8 | 4.44 | 20.1 |
| M14 | 1.81 | >100 | 0.93 | >100 | 5.53 | 37.1 | 4.13 | 19.4 |
| MDA-MB-435 | 0.39 | 1.79 | 0.23 | 0.60 | 7.58 | 67.8 | 1.85 | 4.02 |
| SK-MEL-2 | 2.23 |  | 1.18 | 84.6 | 70.3 | >100 | 4.41 | 21.3 |
| SK-MEL-28 | 5.17 | >100 | 3.86 | 51.7 | 11.9 | 27.9 | 9.81 | 32.2 |
| SK-MEL-5 | 2.75 |  | 2.60 | 14.8 | 13.6 | >100 | 3.57 | 14.7 |
| UACC-257 | 4.99 | >100 | 21.8 | >100 | 83.6 | >100 | 16.9 | 49.3 |
| UACC-62 | 3.52 | >100 | 0.97 | 33.5 | 10.3 | 46.0 | 8.01 | 37.6 |
| **Ovarian Cancer** |  |  |  |  |  |  |  |  |
| IGROV1 | 2.81 | >100 | 1.62 | 34.2 | 11.8 | 60.8 | 8.44 | 36.7 |
| OVCAR-3 | 2.84 | >100 | 2.09 | >100 | 11.3 | 36.4 | 4.04 | 17.2 |
| OVCAR-4 | 16.1 | >100 | 12.7 | >100 | 2.34 | 6.06 | 15.0 | 61.0 |
| OVCAR-5 | 4.07 | >100 | 3.77 | 56.0 | 5.12 | 26.5 | 11.1 | 28.1 |
| OVCAR-8 | 4.17 | >100 | 4.12 | >100 | 6.28 | 34.0 | 11.7 | 70.5 |
| NCI/ADR-RES | 1.26 | 4.40 | 0.51 | 4.60 | 3.07 | 10.7 | 3.12 | 15.2 |
| SK-OV-3 | 4.47 | >100 | 3.92 | >100 | 51.0 | >100 | 10.8 | 43.0 |
| **Renal Cancer** |  |  |  |  |  |  |  |  |
| 786-0 | 2.06 | >100 | 24.4 | >100 | 2.73 | 6.05 | 15.4 | 31.9 |
| A498 | 97.2 | >100 | 35.0 | >100 | 27.2 | 75.1 | 15.2 | 40.8 |
| ACHN | 6.06 | >100 | 5.47 | 48.2 | 1.95 | 4.71 | 7.14 | 22.2 |
| CAKI-1 | 2.82 | >100 | 1.55 | 72.9 | 3.46 | 21.0 | 3.03 | 16.4 |
| RXF 393 | 3.87 | 76.6 | 3.47 | 43.3 | 4.67 | 17.6 | 8.29 | 22.8 |
| SN12C | 3.77 | >100 | 3.93 | >100 | 16.6 | 94.8 | 15.2 | 46.0 |
| TK-10 | 8.62 | >100 | 15.7 | >100 | 20.4 | 55.0 | 20.7 | 43.5 |
| UO-31 | 2.60 | >100 | 1.65 | >100 | 1.72 | 9.08 | 2.71 | 20.3 |
| **Prostate Cancer** |  |  |  |  |  |  |  |  |
| PC-3 | 3.76 | >100 | 4.01 | >100 | 4.56 | 23.1 | 5.85 | 39.0 |
| DU-145 | 4.25 | >100 | 4.33 | >100 | 7.04 | >100 | 11.8 | 40.2 |
| **Breast Cancer** |  |  |  |  |  |  |  |  |
| MCF7 | 0.52 | 10.0 | 0.29 | 19.3 | 0.39 | 15.2 | 0.59 | 18.3 |
| MDA-MB30.2-231/ATCC | 2.10 | 15.6 | 1.59 | 27.4 | 1.79 | 11.3 | 2.71 | 17.6 |
| HS 578T | 2.20 | 21.0 | 3.80 | >100 | 3.88 | 47.2 | 4.14 | 30.2 |
| BT-549 | 2.96 | >100 | 4.40 | 53.0 | 18.1 | 99.0 | 15.2 | 50.0 |
| T-47D | n.t. | n.t. | n.t. | n.t. | 2.42 | >100 | 3.33 | 83.2 |
| MDA-MB-468 | 3.38 | >100 | 2.82 | >100 | 3.57 | 18.3 | 2.60 | 9.88 |

<sup>a</sup> Data obtained from the NCI in vitro disease-oriented human tumor cell line screen. <sup>b</sup> GI<sub>50</sub>: concentration that inhibit 50% net cell growth. <sup>c</sup> TGI total growth inhibition.

To broaden the investigation of the antiproliferative effects of compounds **10a-t** to cancer cells not covered in the NCI panel and considering the encouraging activity reported for imidazothiadiazole derivatives in previous studies targeting pancreatic cancer, we conducted additional experiments to evaluate the cytotoxicity of these compounds on a panel of cells representative of the subtypes of PDAC. Among the pancreatic cancer cells used in the study BxPC-3, CAPAN-1, and SUIT-2 can be categorized as epithelial subtypes, while PANC-1 and PATU-T are representative of the mesenchymal subtype. Additional experiments were performed on a gemcitabine-resistant clone of PANC-1 (PANC-1 GR [41]) and on the primary cell culture PDAC-3 (obtained from a surgically resected patient, as described previously [42]).

A preliminary screening was conducted using three concentrations (0.1, 1, and 10 µM). Subsequently, the most promising compounds were subjected to cytotoxicity testing at various concentrations (ranging from 312 nM to 40 µM) to determine their IC_50_ values. Notably, compounds **10f**, **10k**, **10l**, and **10q** demonstrated significant antiproliferative activity across all the preclinical models. Particularly, the imidazothiadiazoles **10k** and **10l** exhibited IC_50_ values ranging from 1.04 to 6.90 µM, as shown in Table 3.

**Table 3.** Antiproliferative activity of compounds (Cpd) 10f, 10k, 10l and 10q on pancreatic cancer cells.

| IC<sub>50</sub><sup>a</sup> (μM) ± SEM<sup>b</sup> | | | | | | | |
| --- | --- | --- | --- | --- | --- | --- | --- |
| Cpd | CAPAN-1 | PANC-1 | PATU-T | SUIT-2 | BxPC-3 | PDAC-3 | PANC-1 GR |
| **10f** | 4.78±0.01 | 7.40±0.13 | 8.43±0.08 | 10.8±0.12 | 16.62±0.26 | >40 | 15.49±2.9 |
| **10k** | 2.64±0.06 | 3.16±0.007 | 3.73±0.12 | 2.19±0.17 | 2.67±0.15 | 5.54±0.33 | 6.90±0.05 |
| **10l** | 1.54±0.04 | 1.57±0.08 | 1.56±0.007 | 1.70±0.14 | 1.04±0.08 | 2.98±0.4 | 3.44±0.26 |
| **10q** | 6.38±0.1 | 8.65±0.03 | 12.68±0.7 | 6.6±0.1 | 8.83±0.64 | 12.94±2.7 | 15.92±2.6 |

<sup>a</sup> The values are means ± SEM of three separate experiments.
<sup>b</sup> SEM: Standard Error of the Mean.

Finally, we conducted additional experiments to assess the *in vitro* cytotoxicity of the new compound **10l** against normal fibroblasts (Hs27). The outcomes of these experiments enabled us to calculate the selectivity index (SI, calculated as IC_50_ in non-tumor cell line/IC_50_ in the tumor cells PDAC-3), resulting in values of 8.1 for compound **10l**, 5.6 for gemcitabine and 2.6 for defactinib, respectively (Table 4). Consequently, our compounds demonstrated a selectivity index similar to gemcitabine and significantly superior to defactinib, supporting their high cancer-selective profile and prompting further *in vivo* studies.

**Table 4.** Antiproliferative activity of compound **10l** and of reference compounds (gemcitabine and defactinib) on PDAC-3 tumor cells and Hs27 normal fibroblast cells, reported as 50% inhibitory concentration of cell growth (IC_50_) mean values ± standard error of the mean (SEM) values of three separate experiments.

| Cells | **10l** | Gemcitabine | Defactinib |
| --- | --- | --- | --- |
| **PDAC-3** | **2.9 ± 0.4** | **0.03 ± 0.01** | **0.44 ± 0.06** |
| **Hs27** | **23.6 ± 2.5** | **0.17 ± 0.03** | **1.17 ± 0.27** |
| **SI** | **8.1** | **5.6** | **2.6** |

Abbreviations. SI, Selectivity index (IC<sub>50</sub> non-tumor cell line/IC<sub>50</sub> tumor cell line)

### Inhibition of cell migration

Since the high metastatic potential of PDAC is considered one of the main causes of the poor outcome and the high aggressiveness of this disease, the ability of the compounds **10k** and **10l** in inhibiting cell migration of PDAC cells was investigated by scratch wound-healing assays. These experiments were performed on all preclinical models that were previously studied, including BxPC-3, PATU-T, SUIT-2, CAPAN-1, PANC-1, PANC-1 GR, and PDAC-3 cells.

Briefly, 3 × 10^4^ to 5 × 10^4^ cells/well were seeded into 96-well flat-bottom plates in a volume of 100 µL and incubated for 24 h to create a monolayer. The scratches were created in the center of the wells by scraping with a specific tool equipped with needles. The cells were then treated with the compounds at 5 times the IC_50_ concentrations. The selected concentrations were determined through preliminary experiments, which showed no necrotic effects after 24 hours of exposure. The wound closure was monitored by phase-contrast microscopy and the pictures were captured immediately after scratch (T = 0), and at 8-, 20- and 24-hours post-treatment.

As depicted in Figure 2, compounds **10k** and **10l** exhibited potent inhibition of cell migration, demonstrating consistent activity across all investigated PDAC cells. After 8 hours of exposure, the treated cells already displayed a wider scratch area compared to the control group. Remarkably, after 24 hours, a significant two-fold reduction in migration was observed in all PDAC cell lines treated with compound **10l.** Statistical analyses confirmed the significance of these differences when compared to their respective control groups (i.e., untreated cells) in all cell lines.

**Figure 2.**
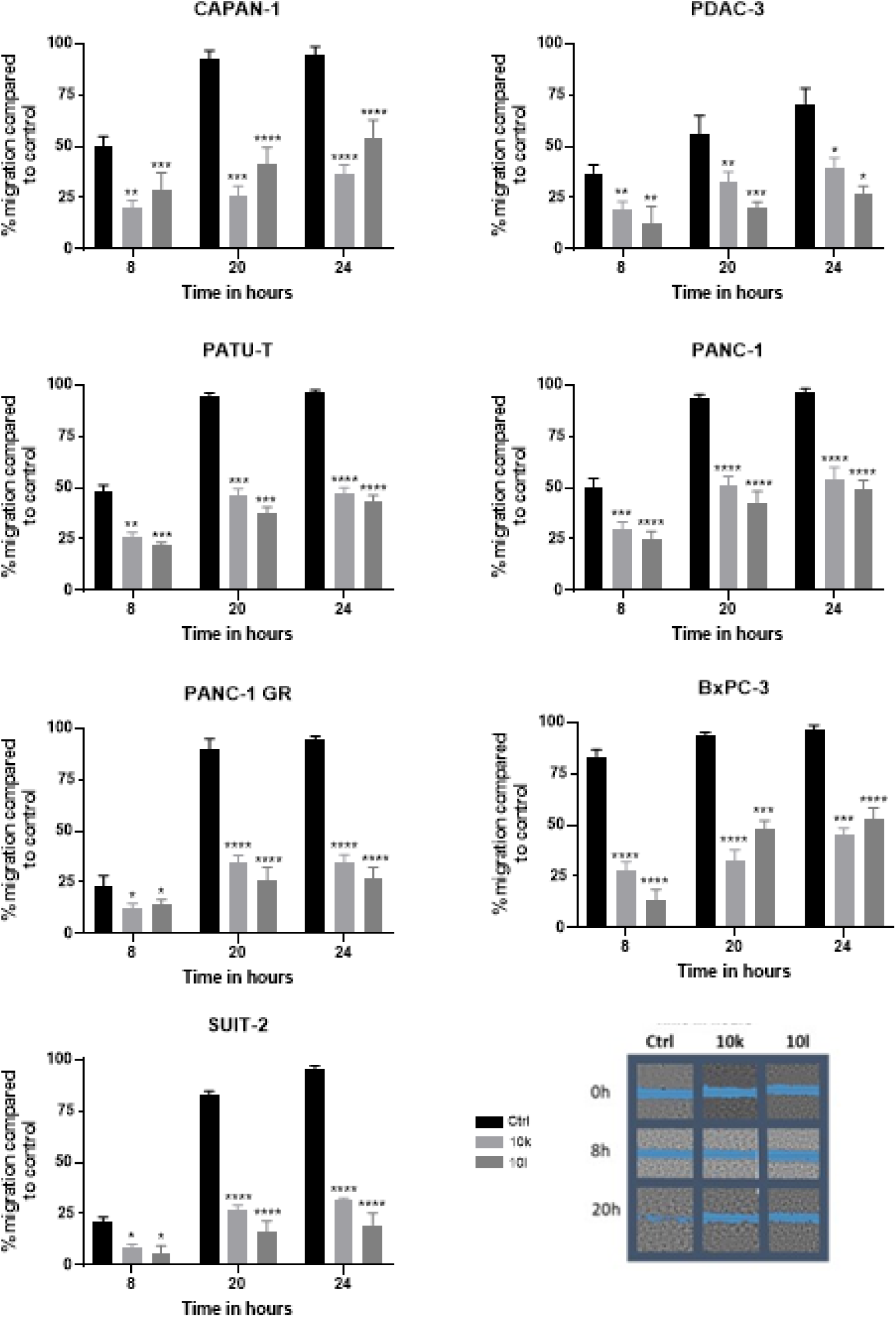
Percentage of migration monitored over time (0, 4, 8, 20 and 24 h) of PANC-1, BxPC-3, PATU-T, CAPAN-1, PDAC-3, PANC-1 GR and SUIT-2 and cells untreated (control, Ctrl) and treated with the compounds **10k** and **10l**. The P values were calculated with Student’s t-test comparing untreated cells with cells treated at concentrations of 5 times IC_50_. ****p < 0.0001, ***p < 0.001, **p < 0.01, *p < 0.05, ns= not significant.

Specifically, when compared to the untreated cells set at 100%, the migration percentages for cells treated with compound **10k** and **10l** were 31% and 19%, respectively, in SUIT-2 cells, 34% and 27%, respectively, in the resistant clone PANC-1 GR, and 39% and 27%, respectively, in PDAC-3 primary cells.

The observed effects on cell migration are consistent with our prior studies on a series of imidazo [2,1-*b*][1,3,4]thiadiazole derivatives, which demonstrated their ability of modulate key regulators of epithelial-to-mesenchymal transition (EMT), including E-cadherin and vimentin, and inhibition of metalloproteinase-2/-9 [12].

### Profiling of inhibition of kinase activity

In order to understand the mechanism of action underlying the above-mentioned promising antiproliferative and antimigratory effects against PDAC cells, we conducted a comprehensive analysis using the Pamgene tyrosine kinase peptide substrate array (PamChip).

This high-throughput assay involves four identical arrays, each comprising 144 peptide sequences immobilized on a porous ceramic membrane. These sequences contain one or more phosphorylation sites, which are derived from literature or computational predictions. The amount of phosphorylated protein by tyrosine kinases from our samples was detected by using specific fluorescently labelled anti-phospho antibodies.

SUIT-2 and PDAC-3 cells, representing the most responsive and resistant cell models, were treated with compound **10l** at a concentration of 5 times the IC_50_ for a duration of 2 hours. Subsequently, both treated and untreated cell lysates were passed through the porous array in the presence of ATP to enable the phosphorylation of peptides by the protein kinases. The assay mixture in both arrays included a fluorescein isothiocyanate (FITC)-labeled antibody, allowing for the quantification of peptide phosphorylation during the incubation period. To capture the real-time reaction kinetics, images measuring the fluorescent signal intensity were taken using a charge-coupled device camera. These kinetic images were captured at 6-second intervals throughout the entire duration of the program.

In SUIT-2 cells, compound **10l** demonstrated significant inhibition of the phosphorylation of 49 protein kinase peptide substrates, with inhibition percentages ranging from 27% to 66% compared to the untreated control. Similarly, in primary PDAC-3 cells, compound **10l** exhibited inhibitory effects on the phosphorylation of 53 different peptide substrates, with inhibition percentages ranging from 26% to 71% compared to the control. Figure 3 illustrates the kinases that showed inhibition percentages exceeding 50% in both cell lines. Remarkably, compound **10l** demonstrated the most remarkable activity against FAK1 and FAK2 in both SUIT-2 and PDAC-3 cell lines, with inhibition percentages of 66% and 71%, respectively.

**Figure 3.**
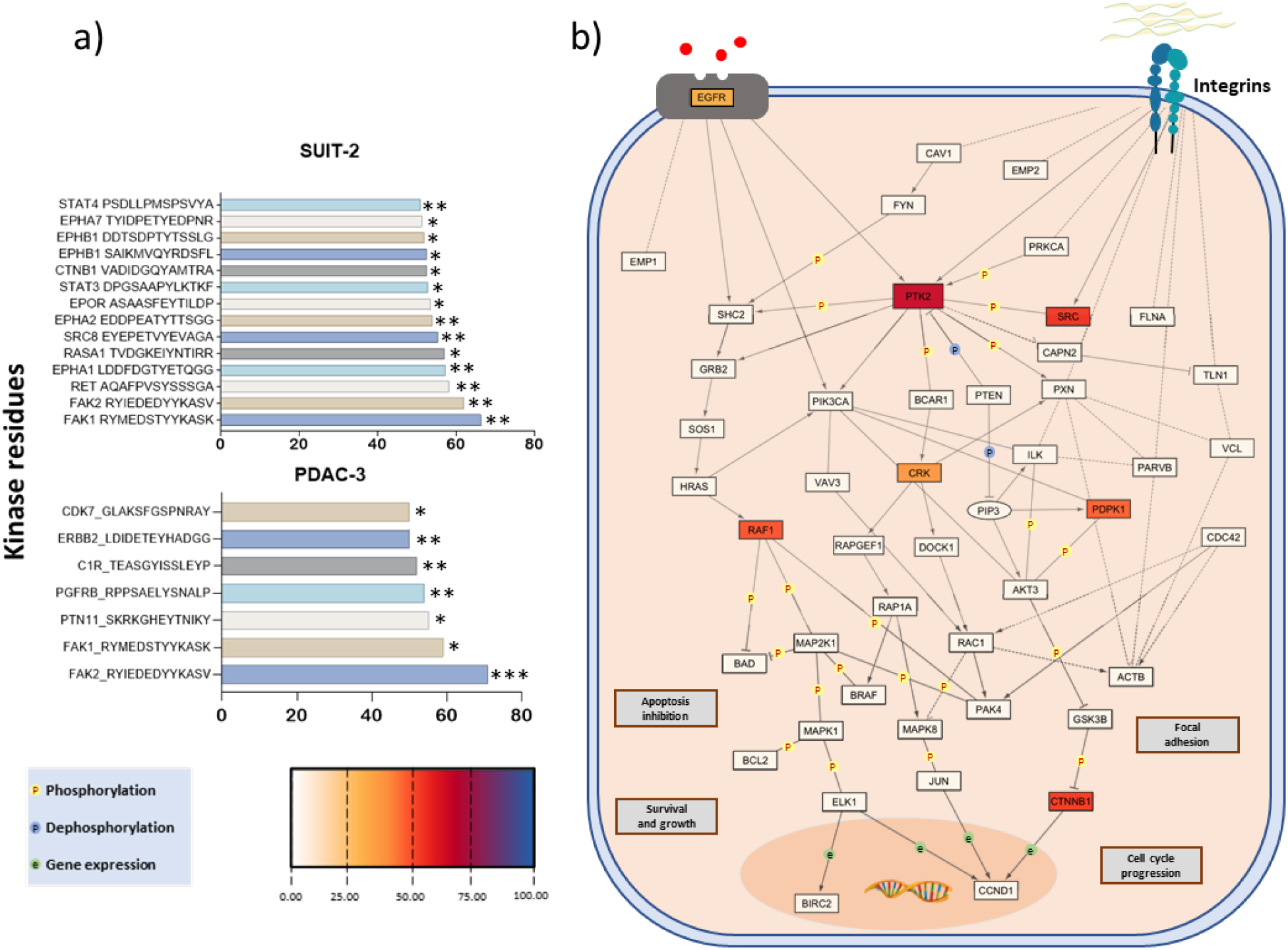
a) Barplot of kinase inhibition in untread cells compared to cells treated with 5 times IC_50_ of **10l**. P values were calculated with Student’s t-test. ***p < 0.001, **p < 0.01, *p < 0.05. b) FAK interaction network obtained with Cytoscape/KEGGscape; the color legend indicates the significance level of each inhibited protein by **10l.**

It is worth mentioning that FAK exhibits known interactions with several kinases inhibited by the compound, including RAF, RAS1, EPHA1, EPHA2, SRC, KDR, and CTNB1, as depicted in Figure 5. For instance, FAK activation can trigger RAS1 activation, facilitating downstream signaling via the MAPK pathway and influencing cellular processes such as cell growth and survival [43]. Moreover, FAK can form complexes with RAF, particularly in integrin-mediated signaling, leading to the activation of the RAF-MEK-ERK cascade, which impacts cell proliferation, survival, and migration [44]. FAK has also demonstrated interactions with Eph receptor family members, such as EPHA1, EPHA2, EPHA7, and EPHB1, modulating cell adhesion and migration and suggesting potential shared downstream signaling pathways [45]. Furthermore, FAK and SRC can establish a reciprocal activation complex through phosphorylation, which plays a critical role in various cellular processes, including cell migration, invasion, and proliferation [46]. In addition, FAK can regulate the phosphorylation and activation of STAT3, contributing to cell survival, proliferation, and migration by activating STAT3-dependent gene expression, as previously described in MIA-PaCa-2 PDAC cancer cells [47]. Lastly, FAK and JAK1 can interact and mutually regulate each other’s activity, as assessed in PDAC preclinical models. In addition, FAK phosphorylation can activate JAK1, initiating downstream signaling events involved in immune response and cytokine signaling [48].

Hence, the obtained results indicatethat the inhibition of FAK phosphorylation serves as a mechanism of action for the new imidazothiadiazole compounds. To validate this mechanism, a specific ELISA assay was employed, and the details of this assay are described in the following section.

### Inhibition of FAK as assessed by ELISA

Considering the intriguing biological activity observed for the new compounds and taking into account the FAK inhibitory potential demonstrated in our previously synthesized series of imidazo [2,1-*b*][1,3,4]thiadiazole derivatives [12], we conducted an analysis using an Enzyme-Linked Immunosorbent Assay (ELISA) to investigate the FAK inhibitory activity of the most active compounds of our new series. Figure 4 illustrates the ability of compounds **10l** and **10k** to inhibit FAK phosphorylation at tyrosine residue 397 (FAK [pY397]), which is crucial for the kinase activity of this protein. Comparatively, both compounds exhibited stronger inhibitory activity than the well-known FAK inhibitor defactinib, used as a reference compound. Specifically, **10l** and **10k** reduced FAK phosphorylation to less than 30% compared to the levels observed in the untreated (control) SUIT-2 and PDAC-3 cells.

**Figure 4.**
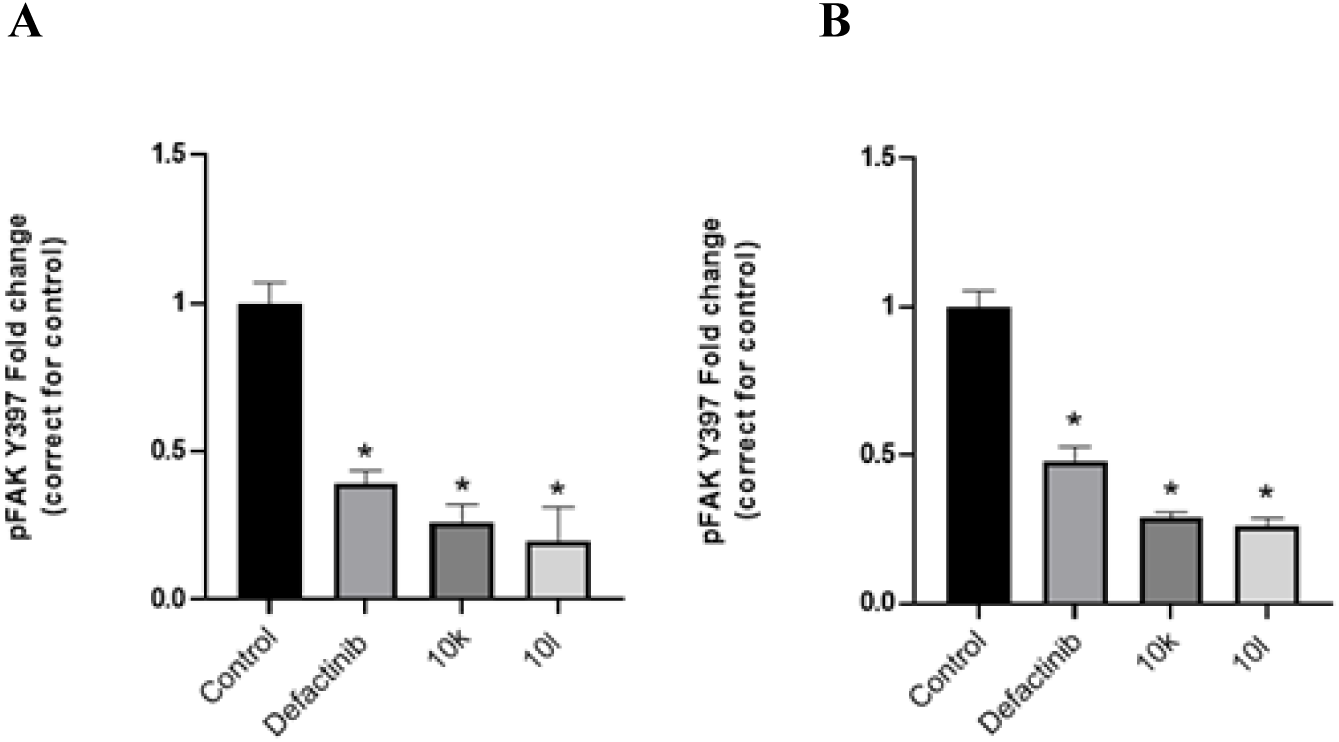
The impact of compounds **10k** and **10l** on the modulation of phosphorylated-FAK at tyrosine residue 397 (pFAK Y397) was assessed in SUIT-2 **(A)** and PDAC-3 **(B)** cells. After treating the cells with compounds **10k**, **10l** at 5 times the IC_50_ concentration for 24 hours, the amount of pFAK was measured in cell lysates. Defactinib, at 5 times IC_50_, was used as a reference compound. The statistical analysis was performed using Student’s t-test, and *p<0.05 denotes significance.

These results are extremely important because FAK is a protein that plays a crucial role in promoting tumorigenesis and cancer progression in several tumors, including PDAC (Kanteti et al. 2016). Additionally, the translocation of FAK in the nucleus induces the inhibition of p53 activity and its downstream gene transcription [50]. As a result, it confers resistance to various anti-cancer treatments, such as chemotherapy and radiation therapy, resulting in treatment failure. Against this background, inhibiting FAK activity can potentially impede tumor growth and metastasis by disrupting these protumorigenic features.

### Effects on cell-cycle modulation

Because of the key role of FAK in cell cycle regulation by promoting cell proliferation and progression through the G1 phase, we assessed the impact of the most cytotoxic compund (compound **10l**) on cell cycle dynamics of SUIT-2 and PDAC-3 cells after 24-hour exposure. Cell cycle progression was analyzed by cytofluorimetry, using propidium iodide (3,8-diamino-5-[3-(diethylmethylammonio)propyl]-6phenyl-diiodide, PI) staining solution.

In both cell lines, the compound **10l** demonstrated potent activity in arresting cell cycle in phase G2/M. Specifically, after 24 hours of treatment, more than 70 and 90% of total SUIT-2 cells and PDAC-3 cells were observed to be blocked in the G2/M phase, respectively (Figure 5). These findings align with the mechanism of action involving FAK inhibition, which is known to induce cell cycle arrest at the G2/M phase and a significant reduction in the S phase [13].

**Figure 5.**
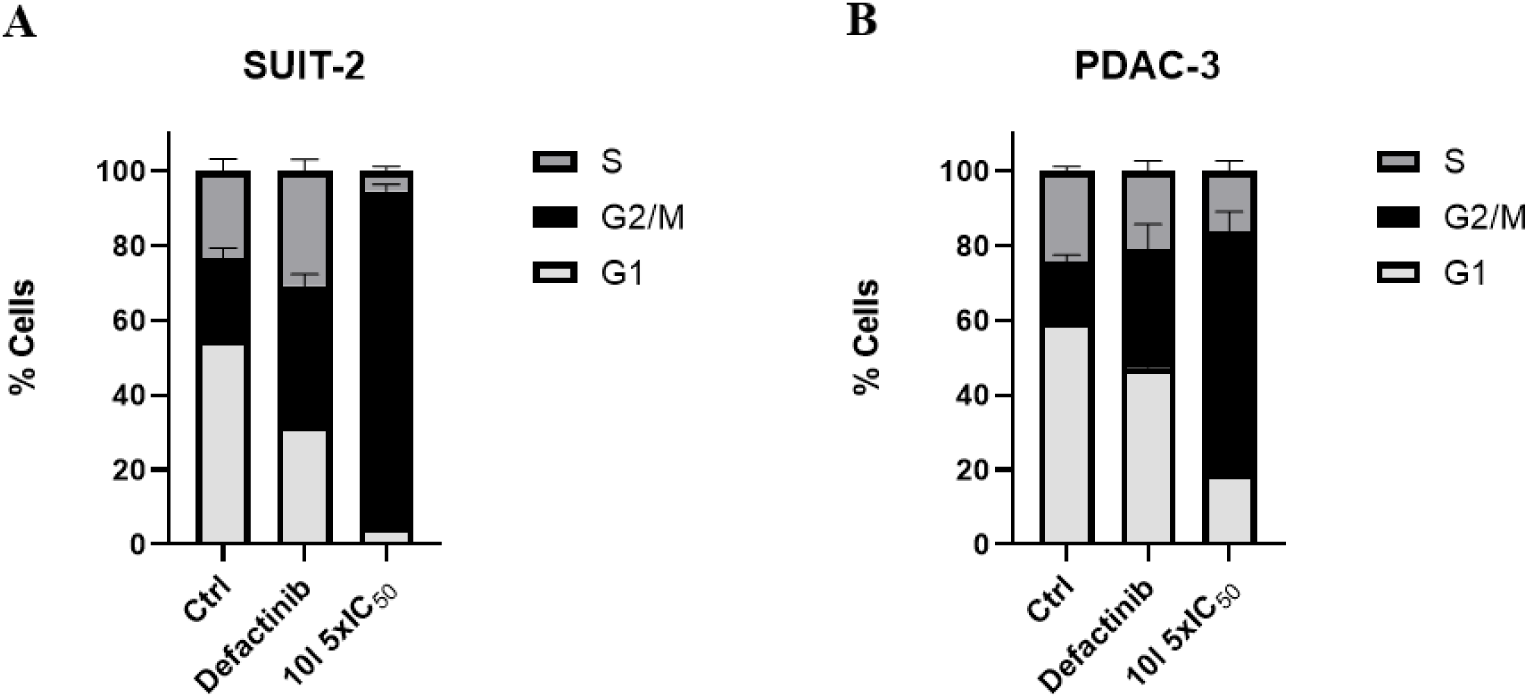
Effects of imidazothiadiazole compound **10l** on cell cycle modulation. SUIT-2 (A) and PDAC-3 (B) cells were treated for 24 hours. Stacked bar graphs show the mean percentage of cells at various stages of cell cycle, G1 (yellow), S (grey), and G2/M (red) phase, in untreated control and after treatment with the compound. Error bars report standard deviations of three separate experiments. Defactinib, at 5-times IC_50_, was used as a reference compound.

### Effects on apoptosis induction

FAK can influence apoptosis signaling pathways, with its overexpression often associated with enhanced cell survival and resistance to apoptosis. Thus, induction of apoptotic response after exposure to compound **10l** was evaluated by the analysis of the stimulation of caspase-3 in both SUIT-2 and PDAC-3 cells (Figure 6).

**Figure 6.**
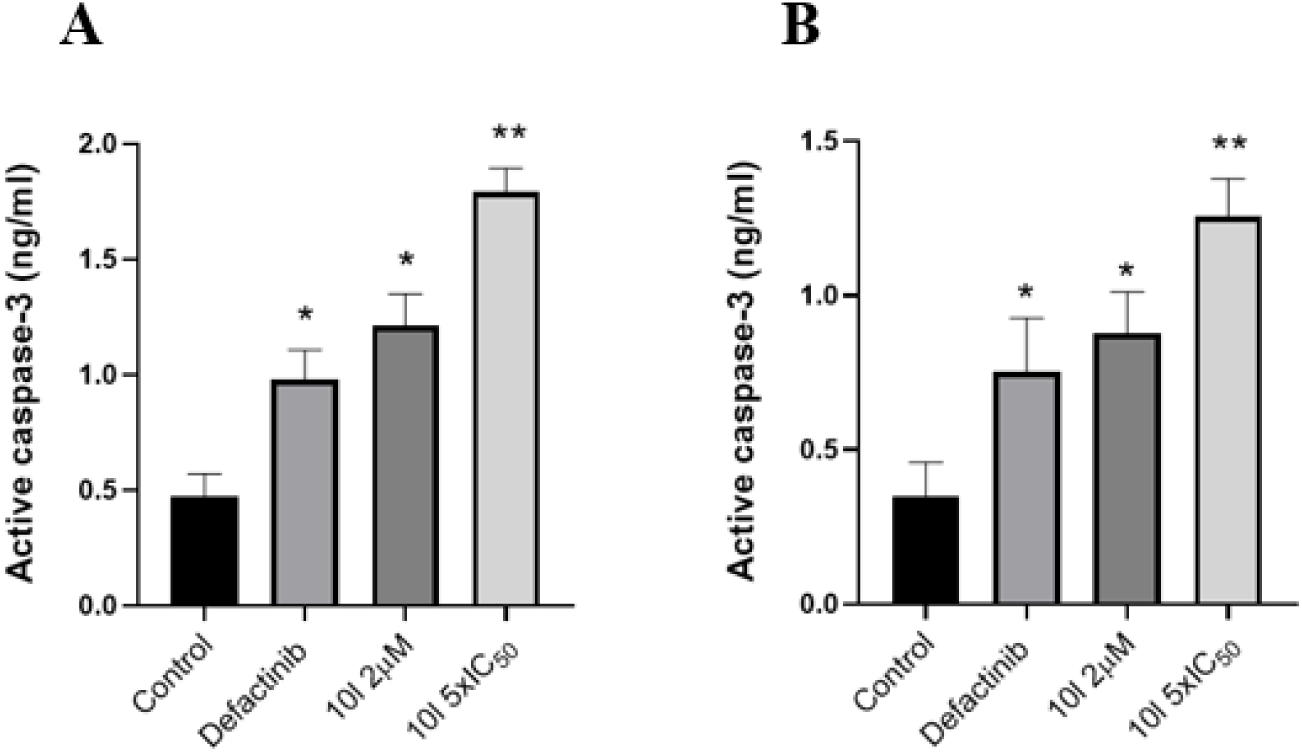
Induction of apoptosis as assessed by evaluation of the levels of active caspase-3 in SUIT-2 cells (A) and PDAC-3 cells (B) treated with **10l**, or defactinib, used at 5 times IC_50_, for 24 hours, compared to control/untreated cells. Measurements were performed in triplicate, and data are presented as means ± SD. *p < 0.01 versus control; **p < 0.001 versus control.

The immunoassay conducted in the present study revealed significantly higher levels of active caspase-3 in SUIT-2 cells that were treated with **10l** and defactinib, compared to the untreated cells. Similar results were observed in PDAC-3 cells, although the levels of active caspase-3 were slightly lower (0.34 ng/mL in untreated cells, 0.75 ng/mL in defactinib-treated cells, and 0.87 ng/mL in **10l**-treated cells). In both PDAC models, the induction of apoptosis was directly proportional to the increasing concentrations of **10l**. Of note, we observed that compound **10l** at 2 μM induces high levels of active Caspase-3 protein expression, achieving a comparable result to defactinib, which was used at 5 times its IC_50_.

The maximum effect was observed in SUIT-2 cells treated with **10l**, resulting in a concentration of active caspase-3 of 1.75 ng/mL.

### Effects on spheroids

FAK plays a role in promoting the attachment of cancer cells to the extracellular matrix, thereby facilitating their invasive potential and aiding in the progression of invasive diseases such as PDAC. Additionally, FAK has been implicated in the formation and maintenance of spheroid structures in PDAC cells, which are associated with increased resistance to therapy and tumor progression [51]. Thus, we conducted further evaluations on three dimensional (3D) spheroids of PDAC-3 cells. These primary cultures successfully formed spheroids that closely resemble the *in vivo* aggregation of tumor cells, consistent with findings from our previous studies [12]. After three days of growth, spheroids were treated with compound **10l** at IC_50_ concentration and concentration 5 times the IC_50_. Additionally, spheroids treated with defactinib and gemcitabine were used as positive control at concentration 5 times IC_50_. As summarized in Figure 7A, both treatment concentrations of compound **10l** inhibit tumor cell proliferation and reduction of the spheroid volume after 11 days. This decrease is plotted as the fold-change between treated and control spheroids and was statistically significant (p-value < 0.0001). A representative picture was captured on Day 1, and subsequent treatments were repeated every four days (Day 5 and Day 8) with corresponding images taken immediately before each treatment (Figure 7B). Moreover, to provides a comprehensive overview of the cell responses to therapy, viability and cell death were analysed by live fluorescence confocal imaging in our 3D models (Figure 7C). Indeed, employing confocal fluorescence microscopy with a correct staining strategy enabled us to visual examination of the inner core of spheroids [52]. After proving antiproliferative activity on 2D cell culture, compound **10l** confirmed its promising activity on 3D PDAC models which better represent the complex tumor structure.

**Figure 7.**
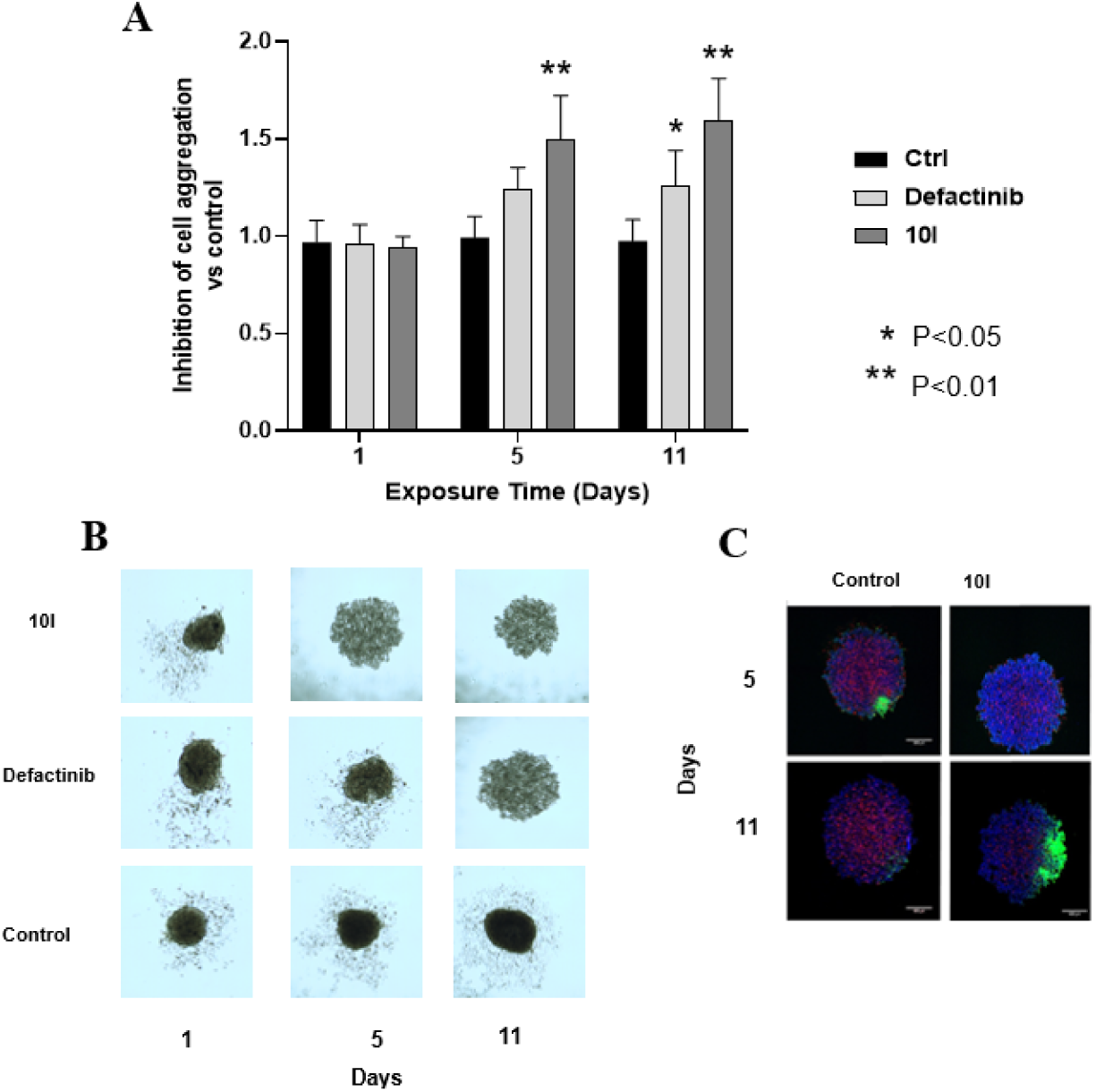
Size reduction of PDAC-3 spheroids treated with compounds **10l** at 5 times the IC_50_ (i.e. 15µM); A) Inhibition of cell aggregation, corrected for control, of the spheroids, at Day 1, Day 5, Day 11; B) Representative pictures of PDAC-3 spheroids exposed to **10l** and gemcitabine, taken at day 1 of treatment, Day 5, and Day 11 with an automated phase-contrast microscope. C) Representative pictures of PDAC-3 spheroids exposed to **10l**, compared to control, taken at day 5 of treatment, and Day 11 with a live fluorescence confocal microscope. p-values were determined by Two-way ANOVA followed by Tukey’s multiple comparisons test, **** = p < 0.0001, ** = p < 0.01. The values were obtained considering the mean values of the areas of at least 8 different spheroids, in two different separated experiments.

### Effects on *in vivo* models

To confirm our findings from *in vitro* PDAC models, we further analysed our most promising compound **10l** on *in vivo* model. Therefore, compound **10l** was tested on a subcutaneous xenograft model, obtained via subcutaneous injection of 1 × 10^6^ primary PDAC-3 cells (Figure 8a).

**Figure 8.**
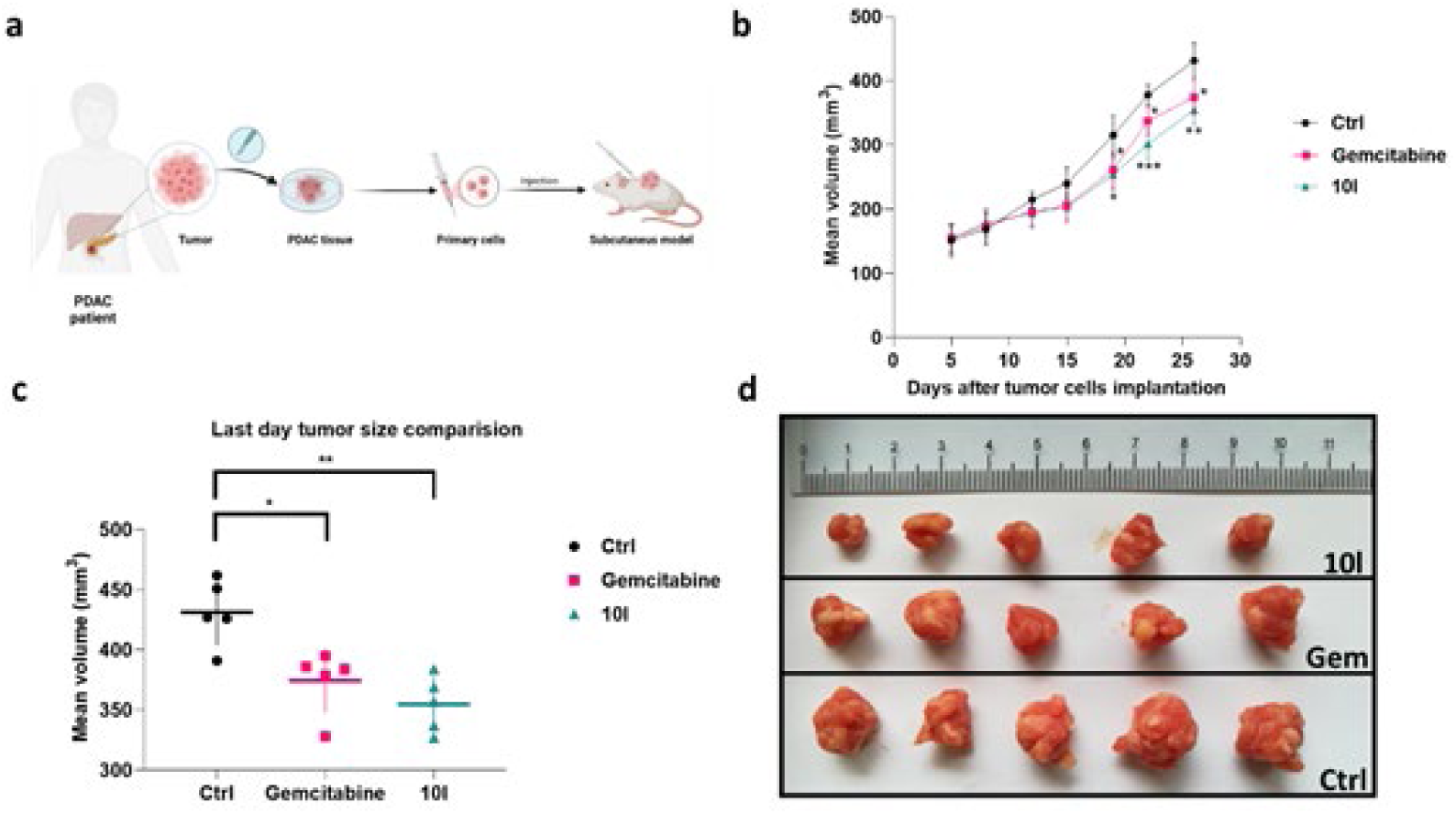
*In vivo* tumor growth results. a) Schematic illustration of tumor inoculation to the obtain subcutaneous model. b) Tumor growth curves show a significate difference between control and treated groups (gemcitabine and **10l**) from day 19th (mean ±SD). c) Significant tumors volume reduction after 2 weeks of treatments d) representative tumors’ images from mice subcutaneously injected with PDAC-3 cells.

After 5 days from tumor implantation, when tumor volume threshold (100 mm^3^) was reached, cohorts of mice were randomised into different treatment groups (N=5 each). Three experimental groups were treated with 1) **10l** (25 mg/kg) three times for two week or 2) gemcitabine (100 mg/kg) or 3) two times per week for two weeks.

Remarkably, compound **10l** demonstrated a significant effect (p < 0.05) on mice already after two weeks from the initial treatment, leading to a reduction in tumor volume compared to the untreated cohort (Figure 8b). After three weeks, all experimental groups exhibited a substantial reduction in tumor volume (Figure 8c,d). Animals treated with **10l** displayed a significant 17.7% reduction in tumor mass compared to the untreated/control group (354.8 vs. 431.4 mm³, p < 0.05). The gemcitabine group showed a similar effect with a 13.2% reduction in tumor mass compared to untreated/control animals (374.4 vs. 431.4 mm³, p < 0.05).

Of note, **10l** was well tolerated; no toxic deaths or signs of toxicity were observed, and the body weight of animals given **10l** was similar to that of the untreated group, as reported in Figure 9. To summarize, our *in vitro* evidence has been confirmed in an *in vivo* model where the administration of compound **10l** demonstrated comparable inhibition of tumor growth to that observed with gemcitabine, which is commonly used in the clinical treatment of PDAC.

**Figure 9.**
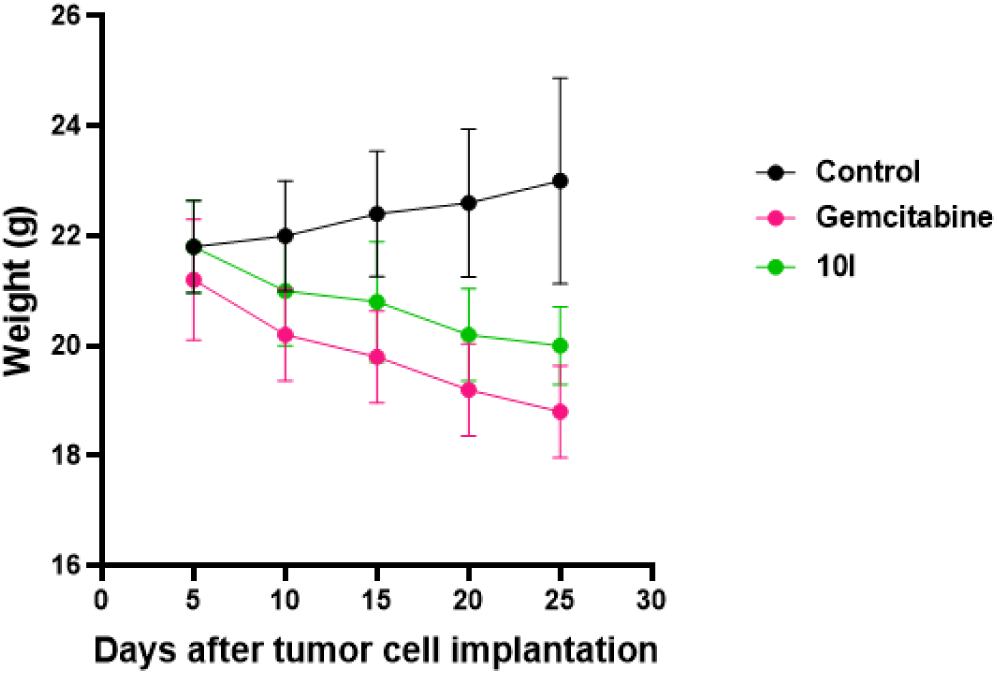
Body weight of mice monitored during treatment.

## Conclusions

FAK is an important emerging target in PDAC because it enhances cancer cell survival by inhibiting apoptosis and prompting cell proliferation signals, while promoting cell migration, immune evasion, and tissue remodeling. In this study, we reported the synthesis and the ability to inhibit cell proliferation and migration in PDAC cells of a novel series of imidazo[2,1-*b*][1,3,4]thiadiazole compounds. Four of the new derivatives, **10f**, **10k**, **10l**, and **10q**, exhibited potent antiproliferative activity against PDAC cell lines showing IC_50_ values ranging from 1.04 to 16.62 µM. High-throughput kinase arrays, carried out for the most promising compound **10l**, unveiled a noteworthy suppression of FAK phosphorylation in SUIT-2 and PDAC-3 cells, suggesting it as the main mecchanism of action. This hypothesis was subsequently confirmed by a specific ELISA assay. The *in vitro* results related to the effects of these compounds on cell migration, as well as their ability to block the cell cycle at the G2/M phase, are congruent with a mechanism involving FAK inhibition. Noteworthy, the significant antiproliferative activity of compound **10l** was corroborated in 3D spheroids of PDAC-3 cells and in a xenograft *in vivo* model, in which a significant reduction of tumor volume was observed after two weeks of treatment. Additionally, this compound did not cause any signs of toxicity and no reduction in mice body weight. This aligns with its low selectivity index value *in vitro*, prompting forthcoming *in vivo* studies to validate this safety profile, which is crucial for the strategic planning of future clinical trials.

Altogether, the obtained results revealed the promising anticancer properties of a new series of FAK inhibitors with imidazothiadiazole structure laying the foundation for the development of new therapeutic approaches in the treatment of PDAC disease.

## Acknowledgments / Funding

The authors thanks A. Griffioen (Angiogenesis-group, Amsterdam UMC) for the migration station to conduct wound-healing assays and the members of the EORTC-PAMM group for the useful discussion and support.

F.S. and E.G. acknowledge EU funding by the H2020-MSCA-RISE program through ALISE (#101007256) project, KWF Dutch Cancer Society, grant number 13598, and Associazione Italiana per la Ricerca sul Cancro (AIRC/Start-Up grant).

## References

[1] Rawla P, Sunkara T, Gaduputi V. Epidemiology of Pancreatic Cancer: Global Trends, Etiology and Risk Factors. World J Oncol. 2019;10(1):10–27. doi: 10.14740/wjon1166.

[2] Siegel RL, Giaquinto AN, Jemal A. Cancer statistics, 2024. CA Cancer J Clin. 2024;74(1):12–49. doi: 10.3322/caac.21820.

[3] Randazzo O, Papini F, Mantini G, Gregori A, et al. «Open Sesame?»: Biomarker Status of the Human Equilibrative Nucleoside Transporter-1 and Molecular Mechanisms Influencing Its Expression and Activity in the Uptake and Cytotoxicity of Gemcitabine in Pancreatic Cancer. Cancers, 2020;12 (11):3206. doi:10.3390/cancers12113206.

[4] Capula M, Macarena P, Geng X, et al. Role of Drug Catabolism, Modulation of Oncogenic Signaling and Tumor Microenvironment in Microbe-Mediated Pancreatic Cancer Chemoresistance. Drug Resist Updat. 2022;64:100864. doi: 10.1016/j.drup.2022.100864.

[5] Bazeed AY, Day CM, Garg S. Pancreatic Cancer: Challenges and Opportunities in Locoregional Therapies. Cancers. 2022;31;14(17):4257. doi: 10.3390/cancers14174257.

[6] Halbrook CJ, Lyssiotis CA, Pasca di Magliano M, et al. Pancreatic cancer: Advances and challenges. Cell. 2023;13;186(8):1729–1754. doi: 10.1016/j.cell.2023.02.014.

[7] Conroy T, Hammel P, Hebbar M, et al. FOLFIRINOX or Gemcitabine as Adjuvant Therapy for Pancreatic Cancer. N Engl J Med. 2018;20;379(25):2395–2406. doi: 10.1056/NEJMoa1809775

[8] Conroy T, Castan F, Lopez A, Turpin A, et al. Five-Year Outcomes of FOLFIRINOX vs Gemcitabine as Adjuvant Therapy for Pancreatic Cancer: A Randomized Clinical Trial. JAMA Oncol. 2022;1;8(11):1571–1578. doi: 10.1001/jamaoncol.2022.3829.

[9] Nitipir C, Vrabie R, Parosanu AI, et al. Clinical Impact of the Administration of FOLFIRINOX Beyond Six Months in Advanced Pancreatic Adenocarcinoma: A Cohort Study. Cureus. 2021;8;13(11):e19361. doi: 10.7759/cureus.19361.

[10] Puik JR, Swijnenburg RJ, Kazemier G, et al. Novel Strategies to Address Critical Challenges in Pancreatic Cancer. Cancers (Basel). 2022;25;14(17):4115. doi: 10.3390/cancers14174115.

[11] Kung HC, Yu J. Targeted therapy for pancreatic ductal adenocarcinoma: Mechanisms and clinical study. MedComm (2020). 2023;19;4(2):e216. doi: 10.1002/mco2.216.

[12] Li Petri G, Cascioferro S, El Hassouni B, et al. Biological Evaluation of the Antiproliferative and Anti-migratory Activity of a Series of 3-(6-Phenylimidazo[2,1-b][1,3,4]thiadiazol-2-yl)-1H-indole Derivatives Against Pancreatic Cancer Cells. Anticancer Res. 2019;39(7):3615–3620. doi: 10.21873/anticanres.13509.

[13] Mustafa M, Abd El-Hafeez AA, Abdelhafeez DA. FAK Inhibitors as Promising Anticancer Targets: Present and Future Directions. Fut Med Chem. 2021;13(18):1559–1590. doi:10.4155/fmc-2021-0015.

[14] Cance WG, Kurenova E, Marlowe T, et al. Disrupting the scaffold to improve focal adhesion kinase-targeted cancer therapeutics. Sci Signal. 2013;26;6(268):pe10. doi: 10.1126/scisignal.2004021.

[15] Mitra SK, Hanson DA, Schlaepfer DD. Focal adhesion kinase: in command and control of cell motility. Nat Rev Mol Cell Biol. 2005;6(1):56–68. doi: 10.1038/nrm1549.

[16] Kurenova E, Li-Hui X, Xihui Y. Focal Adhesion Kinase Suppresses Apoptosis by Binding to the Death Domain of Receptor-Interacting Protein. Mol Cell Biol. 2004;(10):4361–4371. doi:10.1128/MCB.24.10.4361-4371.2004.

[17] Golubovskaya VM, Cance WG. FAK and p53 protein interactions. Anticancer Agents Med Chem. 2011;11(7):617–9. doi: 10.2174/187152011796817619.

[18] Nguyen K, Yan Y, Yuan B, Dasgupta A, et al. ST8SIA1 Regulates Tumor Growth and Metastasis in TNBC by Activating the FAK–AKT–mTOR Signaling Pathway. Mol Cancer Ther. 2018;17(12):2689–2701. doi:10.1158/1535-7163.MCT-18-0399.

[19] Chauhan A, Tabassum K. Focal adhesion kinase-An emerging viable target in cancer and development of focal adhesion kinase inhibitors. Chem Bio Drug Design 2021;97(3): 774–794. doi:10.1111/cbdd.13808.

[20] Sun J, Yang YS, Li W, et al. Synthesis, biological evaluation and molecular docking studies of 1,3,4-thiadiazole derivatives containing 1,4-benzodioxan as potential antitumor agents. Bioorg Med Chem Lett. 2011;15;21(20):6116–21. doi: 10.1016/j.bmcl.2011.08.039.

[21] Duan YT, Yao YF, Huang W, et al. Synthesis, biological evaluation, and molecular docking studies of novel 2-styryl-5-nitroimidazole derivatives containing 1,4-benzodioxan moiety as FAK inhibitors with anticancer activity. Bioorg Med Chem. 2014;22(11):2947–54. doi: 10.1016/j.bmc.2014.04.005.

[22] Bolchi C, Bavo F, Appiani R, et al. 1,4-Benzodioxane, an evergreen, versatile scaffold in medicinal chemistry: A review of its recent applications in drug design. Eur J Med Chem. 2020;200:112419. doi: 10.1016/j.ejmech.2020.112419.

[23] Elbadawi MM, Eldehna WM, Abd El-Hafeez AA, et al. 2-Arylquinolines as novel anticancer agents with dual EGFR/FAK kinase inhibitory activity: synthesis, biological evaluation, and molecular modelling insights. J Enzyme Inhib Med Chem. 2022;37(1):349–372. doi: 10.1080/14756366.2021.2015344.

[24] Tsai PC, Chu CL, Fu YS, et al. Naphtho[1,2-b]furan-4,5-dione inhibits MDA-MB-231 cell migration and invasion by suppressing Src-mediated signaling pathways. Mol Cell Biochem. 2014;387(1-2):101–11. doi: 10.1007/s11010-013-1875-4.

[25] Carbone D, Vestuto V, Ferraro MR, et al. Metabolomics-assisted discovery of a new anticancer GLS-1 inhibitor chemotype from a nortopsentin-inspired library: From phenotype screening to target identification. Eur J Med Chem. 2022;234:114233. doi: 10.1016/j.ejmech.2022.114233.

[26] Carbone D, Pecoraro C, Panzeca G, et al. 1,3,4-Oxadiazole and 1,3,4-Thiadiazole Nortopsentin Derivatives against Pancreatic Ductal Adenocarcinoma: Synthesis, Cytotoxic Activity, and Inhibition of CDK1. Mar Drugs. 2023;21(7):412. doi: 10.3390/md21070412.

[27] Di Franco S, Parrino B, Gaggianesi M, et al. CHK1 inhibitor sensitizes resistant colorectal cancer stem cells to nortopsentin. iScience. 2021;24(6):102664. doi: 10.1016/j.isci.2021.102664.

[28] Pecoraro C, De Franco M, Carbone D, et al. 1,2,4-Amino-triazine derivatives as pyruvate dehydrogenase kinase inhibitors: Synthesis and pharmacological evaluation. Eur J Med Chem. 2023 Mar 5;249:115134. doi: 10.1016/j.ejmech.2023.115134.

[29] Pecoraro C, Parrino B, Cascioferro S, et al. A New Oxadiazole-Based Topsentin Derivative Modulates Cyclin-Dependent Kinase 1 Expression and Exerts Cytotoxic Effects on Pancreatic Cancer Cells. Molecules. 2021;27(1):19. doi: 10.3390/molecules27010019.

[30] Turdo A, Glaviano A, Pepe G, et al. Nobiletin and Xanthohumol Sensitize Colorectal Cancer Stem Cells to Standard Chemotherapy. Cancers (Basel). 2021;13(16):3927. doi: 10.3390/cancers13163927.

[31] Carbone D, Parrino B, Cascioferro S, et al. 1,2,4-Oxadiazole Topsentin Analogs with Antiproliferative Activity against Pancreatic Cancer Cells, Targeting GSK3β Kinase. ChemMedChem. 2021;16(3):537–554. doi: 10.1002/cmdc.202000752.

[32] Du J, Chu W, Zhang M, Ma C, Feng W. A novel method for preparing Eligulstat through chiral resolution. Bioorg Med Chem Lett. 2020;30(16):127209. doi: 10.1016/j.bmcl.2020.127209.

[33] Vyas VK, Bhalchandra MB. Catalytic Asymmetric Synthesis of β-Triazolyl Amino Alcohols by Asymmetric Transfer Hydrogenation of α-Triazolyl Amino Alkanones. Tetrahedron: Asymmetry 2017;28(7):974–982. doi:10.1016/j.tetasy.2017.05.012.

[34] Maftouh M, Avan A, Funel N, et al. miR-211 modulates gemcitabine activity through downregulation of ribonucleotide reductase and inhibits the invasive behavior of pancreatic cancer cells. Nucleosides Nucleotides Nucleic Acids. 2014;33(4-6):384–93. doi: 10.1080/15257770.2014.891741

[35] Van Der Steen N, Keller K, Dekker H, et al. Crizotinib sensitizes the erlotinib resistant HCC827GR5 cell line by influencing lysosomal function. J Cell Physiol. 2020;235(11):8085–8097. doi: 10.1002/jcp.29463.

[36] Zhang L, Zhao D, Wang Y, et al. Focal adhesion kinase (FAK) inhibitor-defactinib suppresses the malignant progression of human esophageal squamous cell carcinoma (ESCC) cells via effective blockade of PI3K/AKT axis and downstream molecular network. Mol Carcinog. 2021;60(2):113–124. doi: 10.1002/mc.23273.

[37] Macchia M, Barontini S, Bertini S, et al. Design, synthesis, and characterization of the antitumor activity of novel ceramide analogues. J Med Chem. 2001 Nov 8;44(23):3994–4000. doi: 10.1021/jm010947r.

[38] Sciarrillo R, Wojtuszkiewicz A, El Hassouni B, et al. Splicing modulation as novel therapeutic strategy against diffuse malignant peritoneal mesothelioma. EBioMedicine. 2019;39:215–225. doi: 10.1016/j.ebiom.2018.12.025.

[39] Le Large TYS, Bijlsma MF, El Hassouni B, et al. Focal adhesion kinase inhibition synergizes with nab-paclitaxel to target pancreatic ductal adenocarcinoma. J Exp Clin Cancer Res. 2021;40(1):91. doi: 10.1186/s13046-021-01892-z.

[40] Cavazzoni A, La Monica S, Alfieri R, et al. Enhanced efficacy of AKT and FAK kinase combined inhibition in squamous cell lung carcinomas with stable reduction in PTEN. Oncotarget. 2017;8(32):53068–53083. doi: 10.18632/oncotarget.18087.

[41] Le Large TYS, El Hassouni B, Funel N, et al. Proteomic analysis of gemcitabine-resistant pancreatic cancer cells reveals that microtubule-associated protein 2 upregulation associates with taxane treatment. Ther Adv Med Oncol. 2019;11:1758835919841233. doi: 10.1177/1758835919841233.

[42] Rovithi M, Avan A, Funel N, et al. Development of bioluminescent chick chorioallantoic membrane (CAM) models for primary pancreatic cancer cells: a platform for drug testing. Sci Rep. 2017;7:44686. doi: 10.1038/srep44686.

[43] Bahk YY, Cho IH, Kim TS. A cross-talk between oncogenic Ras and tumor suppressor PTEN through FAK Tyr861 phosphorylation in NIH/3T3 mouse embryonic fibroblasts. Biochem Biophys Res Commun. 2008;377(4):1199–204. doi: 10.1016/j.bbrc.2008.10.157.

[44] Delyon J, Servy A, Laugier F, et al. PDE4D promotes FAK-mediated cell invasion in BRAF-mutated melanoma. Oncogene. 2017 Jun 8;36(23):3252–3262. doi: 10.1038/onc.2016.469.

[45] Buckens OJ, El Hassouni B, Giovannetti E, Peters GJ. The role of Eph receptors in cancer and how to target them: novel approaches in cancer treatment. Expert Opin Investig Drugs. 2020;29(6):567–582. doi: 10.1080/13543784.2020.1762566.

[46] Wang S, Englund E, Kjellman P, et al. CCM3 is a gatekeeper in focal adhesions regulating mechanotransduction and YAP/TAZ signalling. Nat Cell Biol. 2021;23(7):758–770. doi: 10.1038/s41556-021-00702-0.

[47] Hsieh YJ, Tseng SP, Kuo YH, et al. Ovatodiolide of Anisomeles indica Exerts the Anticancer Potential on Pancreatic Cancer Cell Lines through STAT3 and NF-κB Regulation. Evid Based Complement Alternat Med. 2016;2016:8680372. doi: 10.1155/2016/8680372.

[48] Macha MA, Rachagani S, Gupta S, et al. Guggulsterone decreases proliferation and metastatic behavior of pancreatic cancer cells by modulating JAK/STAT and Src/FAK signaling. Cancer Lett. 2013;341(2):166–77. doi: 10.1016/j.canlet.2013.07.037.

[49] Kanteti R, Batra SK, Lennon FE, Salgia R. FAK and paxillin, two potential targets in pancreatic cancer. Oncotarget. 2016;7(21):31586–601. doi: 10.18632/oncotarget.8040.

[50] Zhou J, Yi Q, Tang L. The roles of nuclear focal adhesion kinase (FAK) on Cancer: a focused review. J Exp Clin Cancer Res. 2019;38(1):250. doi: 10.1186/s13046-019-1265-1.

[51] Mohamed AA, Thomsen A, Follo M, et al. FAK inhibition radiosensitizes pancreatic ductal adenocarcinoma cells in vitro. Strahlenther Onkol. 2021;197(1):27–38. doi: 10.1007/s00066-020-01666-0.

[52] Sant S, Johnston PA. The production of 3D tumor spheroids for cancer drug discovery. Drug Discov Today Technol. 2017;23:27–36. doi: 10.1016/j.ddtec.2017.03.002.

